# A nutrient bottleneck limits antibiotic efficacy in structured bacterial populations

**DOI:** 10.1101/2025.03.12.642894

**Authors:** Anna M. Hancock, Arabella S. Dill-Macky, Jenna A. Moore, Catherine Day, Mohamed S. Donia, Sujit S. Datta

## Abstract

Antibiotic resistance is a growing global health threat. Therefore, it is critically important to optimize how existing antibiotics act against bacterial infections. Although antibiotic activity is well studied at the single cell level, many infections are caused by spatially structured multicellular populations. In such populations, cellular consumption of scarce nutrients establishes strong spatial variations in their abundance. These nutrient variations have long been hypothesized to help bacterial populations tolerate antibiotics, since single-cell studies show that antibiotic tolerance depends on metabolic activity, and thus, local nutrient availability. Here, we directly test this hypothesis by visualizing cell death in *Escherichia coli* populations with defined structures upon exposure to nutrient (glucose) and antibiotic (fosfomycin). We find that nutrient availability acts as a bottleneck to antibiotic killing, causing death to propagate through the population as a traveling front—a phenomenon predicted over 20 years ago, but never verified until now. By integrating our measurements with biophysical theory and simulations, we establish quantitative principles that explain how collective nutrient consumption can limit the progression of this “death front,” protecting a population from a nominally deadly antibiotic dose. While increasing nutrient supply can overcome this bottleneck, our work reveals that in some cases, excess nutrient can unexpectedly *promote* the regrowth of resistant cells. Altogether, this work provides a key step toward predicting and controlling antibiotic treatment of spatially structured bacterial populations, yielding fundamental biophysical insights into collective behavior and helping to guide strategies for more effective antibiotic stewardship.

---

As the rise of antibiotic resistance outpaces the discovery of new antibiotics [1, 2], there is an urgent need to optimize how existing antibiotics are administered to prolong and enhance their efficacy against bacterial infections. Current understanding of antibiotic activity is largely based on studies of individual cells in homogeneous liquid cultures. While these studies have yielded powerful insights [3–11], antibiotic treatments successful in liquid cultures often fail against natural bacterial populations—which are typically large, spatially structured, multicellular collectives [12–18]. Addressing this disconnect is a critical challenge for biomedical science and industry.

As nutrient molecules diffuse into a structured population, they are consumed by the cells, creating steep gradients from the surface of the population inward (19, Fig. **1**A). As a result, cells near the surface are more metabolically active [10] and therefore more susceptible to many antibiotics. By contrast, inner cells are more dormant and therefore die slower when exposed to antibiotics, a phenomenon known as antibiotic tolerance [7, 20–22]. This inner reservoir of metabolically dormant cells has long been posited to prolong the the survival of bacterial populations in response to administered antibiotics [23–27]. Unfortunately, systematically testing this idea has been challenging [28] due to the physicochemical heterogeneity of natural bacterial populations [29–32] as well as technical limitations in probing their internal dynamics using conventional microscopy techniques [33]. One way to overcome these challenges is to confine cells to concentrated 2D packings using microfluidics, providing useful insights into the response of bacteria to antibiotic and nutrient gradients separately [34–38]. However, studies of how nutrient and antibiotic transport jointly influence cell death—particularly in populations with defined spatial structures and cell concentrations that more closely mimic the real world—are lacking.

**Fig. 1.**
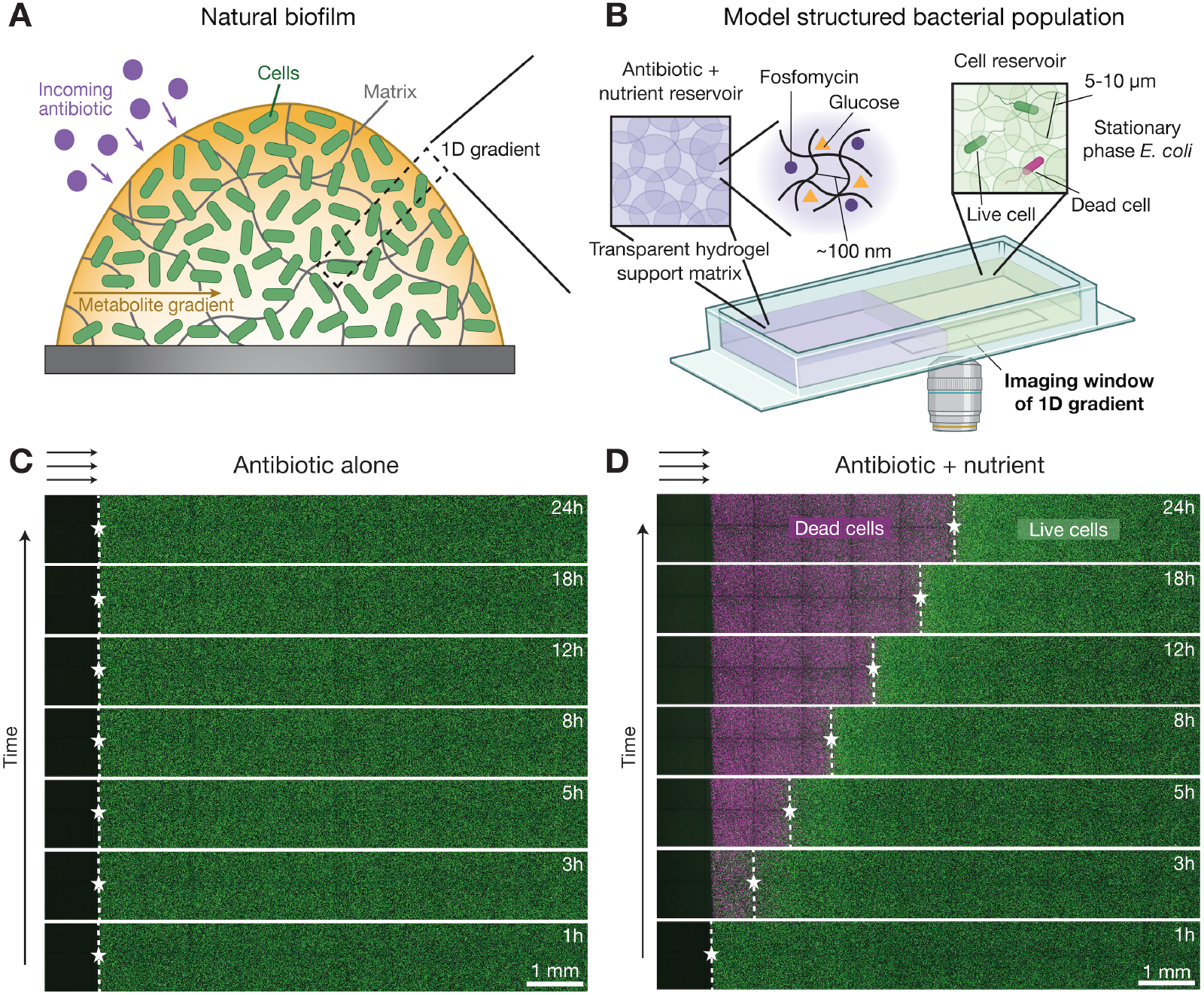
Nutrients unlock a front of cell death that propagates through a structured *E. coli* population exposed to antibiotic. **A** Natural bacterial populations, such as biofilms, are spatially structured, with chemical gradients extending inward from their surfaces. **B** Schematic of our experimental platform, which provides a way to systematically and controllably study such structured populations. We immobilize stationary phase *E. coli* in granular hydrogel matrices (green, right) through which nutrient (glucose) and antibiotic (fosfomycin) can diffuse from a cell-free reservoir (purple, left). The matrices are transparent, enabling direct visualization of cells (green) and their death (magenta) using confocal microscopy. **C** Micrographs showing an *E. coli* population (green, initial concentration *b*_0_ = 10^8^ CFU/mL) encountering fosfomycin (initial concentration *a*_0_ = 2048 µg/mL, equivalent to ∼ 250× MIC) as it diffuses in from the cell-free reservoir on the left (black). Despite the strong antibiotic dose, the cells remain alive over the duration of the experiment, as indicated by the consistent green fluorescence signal from the cells and lack of detectable magenta signal from propidium iodide, a dead cell indicator. **D** Repeating the same experiment, but with 0.22 mM glucose added to the reservoir, reveals a propagating front of cell death, indicated by the dashed lines with stars that mark the replacement of green signal from live cells with magenta signal from dead cells. Micrographs in **C**-**D** are maximum intensity projections of three optical slices taken 50 µm apart starting 50 µm above the bottom of the sample.

Here, we address this gap in knowledge by immobilizing *E. coli* populations of defined structures in transparent hydrogel matrices, enabling direct visualization of cell death upon controlled exposure to both nutrient and antibiotic using fluorescence microscopy. We choose fosfomycin as the test antibiotic because of its common use to treat urinary tract infections, as well as growing interest in using it to treat many other infections [39–41]; moreover, its efficacy is known to be dependent on nutrient availability [42–47], as with many other antibiotics. We use glucose as the nutrient since it is the most bioavailable sugar in the body and does not trigger resistance changes against fosfomycin [43, 48]. This experimental model system reveals that cell death sweeps through a bacterial population as a sharp front whose progression is determined by the coupling between the microscopic chemical processes underlying cell growth and death and the largerscale transport of both nutrient and antibiotic. Building on prior work that first predicted the existence of such death fronts [24, 49], we develop a biophysical framework that elucidates the conditions under which limited nutrient availability acts as a bottleneck to antibiotic efficacy. Our work also uncovers how partial death of a population can enable a resistant subpopulation of cells to scavenge nutrients and regrow in the wake of a death front—a manifestation of heteroresistant population recovery. By shedding new light on the coupling between cellular metabolism and chemical transport in structured bacterial populations, our results could help develop more effective strategies to treat bacteria in health, as well as in agriculture, the environment, and industry more broadly.

## RESULTS

### Cell death propagates through a structured population exposed to antibiotic—but only when nutrient is present

To create bacterial populations with defined spatial structure, we immobilize *E. coli* cell in hydrogel matrices, forming “synthetic biofilms.” Each matrix is made of jammed, biocompatible hydrogel grains swollen in liquid media. The internal mesh size of each grain is ∼100 nm, smaller than the cells, but large enough to allow the transport of oxygen, glucose, and fosfomycin [50–54]. The pores formed in the interstices between grains are ∼ 0.1 − 1 µm in size, tight enough to immobilize each cell in place without impeding its growth [55]. Moreover, because the grains are themselves liquid-infused hydrogels, the matrices are transparent, enabling direct visualization of the cells via confocal microscopy. We use *E. coli* in stationary phase as the initial inoculum to mimic the conditions found within many biofilms [56, 57]. The cells constitutively express green fluorescent protein (GFP) in their cytoplasm; we also mix propidium iodide (PI) into the hydrogel matrix to enable visualization of cell death using fluorescence.

The experimental platform is schematized in Fig. **1**B. To mimic the geometry of natural biofilms, where exogenous nutrient and antibiotic diffuse inward from their surface (Fig. **1**A), we construct each matrix in two sections. One acts as a reservoir containing glucose and fosfomycin at initially defined concentrations *c*_0_ and *a*_0_, but without cells (purple in Fig. **1**B), while the other contains only cells, with no glucose or fosfomycin initially present (green in Fig. **1**B)— representing the exterior and interior of a biofilm, respectively. These two sections are initially separated by an impermeable acrylic partition; at the beginning of each experiment (time *t* = 0), we remove the partition, allowing glucose and fosfomycin to diffuse into the cell-containing section.

We first examine the case of antibiotic exposure under nutrient-free conditions (*c*_0_ = 0). The fosfomycin concentration is *a*_0_ = 2048 µg/mL, over two orders of magnitude larger than the minimum inhibitory concentration (MIC) needed to stop cells from growing in nutrient-rich liquid culture. Surprisingly, despite this large antibiotic concentration, the cells remain alive for the entire duration of the experiment, as indicated by the maintenance of green GFP fluorescence and lack of magenta PI fluorescence in Figs. **1**C (Movie S1) and S1.

Repeating this experiment, but with a small physiological amount [58] of glucose also added to the reservoir (*c*_0_ = 0.22 mM), yields dramatically different results. As shown in Fig. **1**D (Movie S2), a “death front” — indicated by the magenta PI signal — progressively sweeps through the population. This front is remarkably sharp, as indicated by the white dashed lines and stars in Fig. **1**D.

### Death front dynamics are influenced by changes in nutrient availability, but not in antibiotic exposure

How quickly does this death front progress? And what factors control its dynamics? Our findings in Fig. **1**C-D indicate that exposure to antibiotic above MIC is necessary, but not sufficient, to kill cells. Instead, the progression of the death front is constrained by nutrient availability, not antibiotic penetration. To test this idea, we repeat the experiment of Fig. **1**D, but with a ∼ 10-fold reduction in the fosfomycin concentration to *a*_0_ = 256 µg/mL, still over an order of magnitude larger than MIC. We find identical results (Fig. **2**A, purple curve)— further indicating that nutrient availability is the bottleneck to cell killing. In addition, repeating the experiment of Fig. **1**D with a 10-fold increase in the glucose concentration instead yields a markedly faster death front (Fig. **2**B, Movie S3), as expected. This effect is not specific to glucose as the nutrient, but extends to other 6-carbon sugars as well (Fig. S2).

**Fig. 2.**
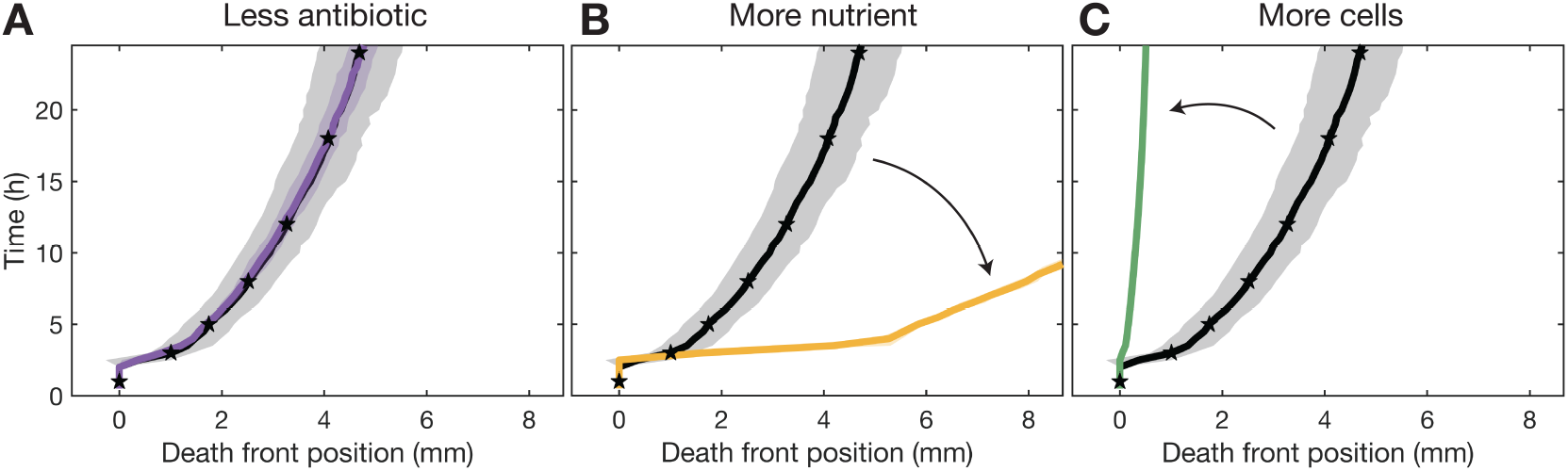
Cellular metabolism of nutrients regulates death front dynamics. **A** Decreasing fosfomycin source concentration from *a*_0_ = 2048 µg/mL (black) to 256 µg/mL (purple) does not change death front propagation. **B** Increasing glucose source concentration from *c*_0_ = 0.22 mM (black) to 2.2 mM (yellow) hastens the death front. **C** Increasing cell concentration from *b*_0_ = 10^8^ CFU/mL (black) to 10^9^ CFU/mL (green) slows the death front. In all three panels, the black line shows the results from the base case of Fig. **1**D (*a*_0_ = 2048 µg/mL, *c*_0_ = 0.22 mM, *b*_0_ = 10^8^ CFU/mL), with stars indicating the same times as in Fig. **1**D and shading indicating standard deviation of results taken across 3 biological replicates.

In addition to diffusing through the hydrogel matrix, the nutrient is actively metabolized by the cells in the population. Thus, we expect that repeating the experiment of Fig. **1**D, but with a 10-fold increase in the concentration of cells, which greatly increases the collective consumption of nutrient, should hinder nutrient availability and hence, the progression of the death front. Our results confirm this expectation, as shown in Fig. **2**C (Movie S4): whereas the initial death front of Fig. **1**D killed half the population after 24 h, in the more concentrated population, less than a tenth of the population is killed in the same duration. These results indicate that because cells must be metabolically active to be killed, nutrient transport and availability acts as a bottleneck to antibiotic efficacy. This finding could help explain why, though inadequate penetration is commonly thought to limit the efficacy of antibiotics against natural bacterial populations, this is often not the case in practice [26, 59].

### A minimal model recapitulates the experimental observations without any fitting parameters

To further rationalize the experimental observations, we build on previous work [24, 60, 61] to construct a continuum model describing the collective dynamics of bacteria, nutrient, and antibiotic, with concentrations *b*(*x, t*), *c*(*x, t*), and *a*(*x, t*), respectively, over a rectilinear domain described by the position coordinate *x*. The model is summarized in Fig. **3**A and detailed in the *SI Appendix*. The entire domain has length *L*, no flux conditions at its boundaries, and is split into two sections, just as in the experiments. At *t* = 0, only nutrient and antibiotic are uniformly distributed in the first section (− *L/*2 ≤ *x* ≤ 0) at concentrations *c*_0_ and *a*_0_, respectively, while only cells are uniformly distributed in the second section (0 ≤ *x* ≤ *L/*2) at a concentration *b*_0_.

**Fig. 3.**
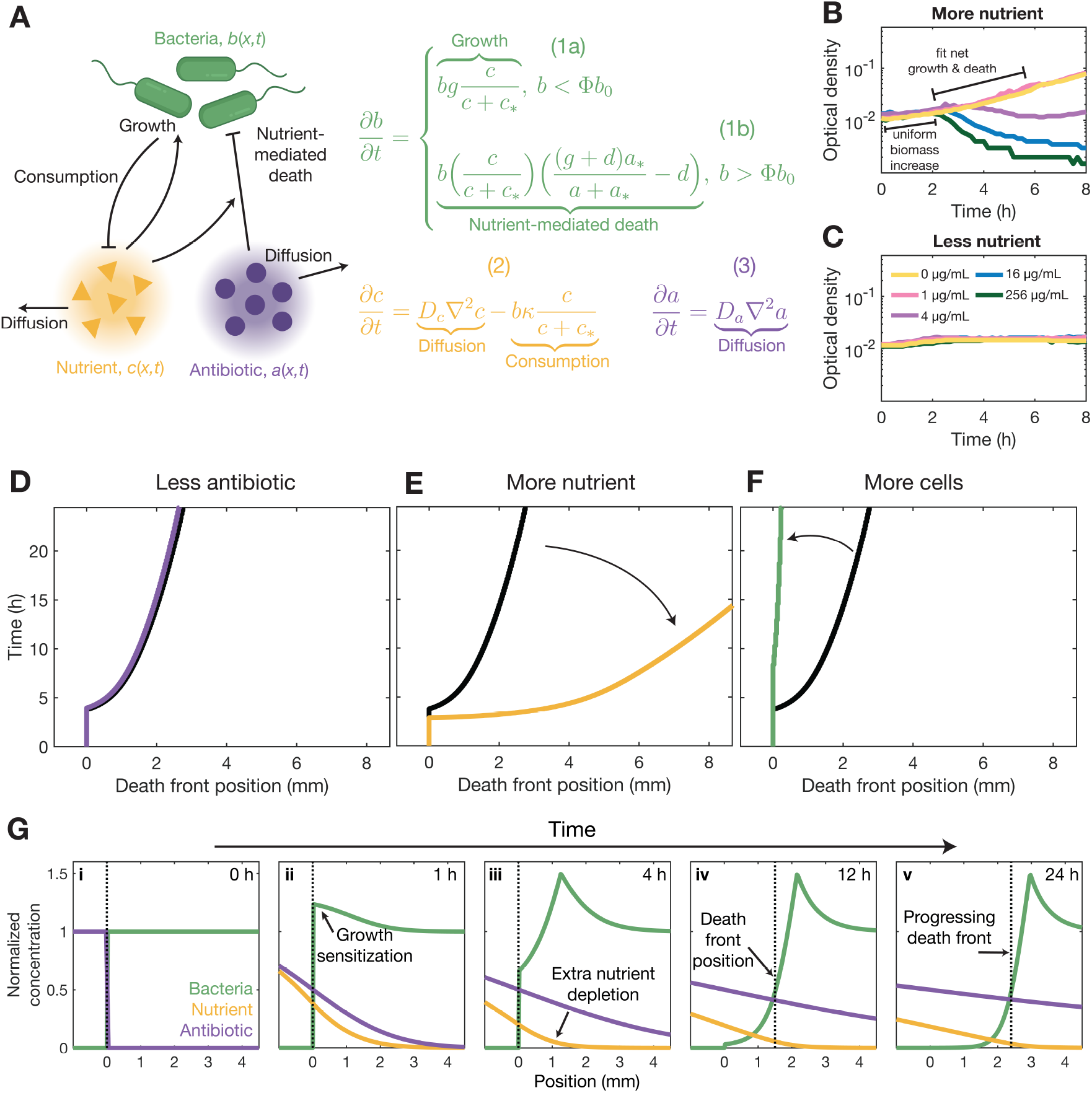
Continuum model of bacteria, nutrient, and antibiotic interactions recapitulates experimental observations of death fronts. **A** The model describes the dynamics of growing and dying bacteria relative to the threshold concentration Φ*b*_0_ (Eq. 1), nutrient that diffuses and is consumed by cells (Eq. 2), and antibiotic that also diffuses (Eq. 3) over an extended one-dimensional domain mimicking the experiments. **B** Measurements of cell growth and death in well-mixed nutrient-replete (*c*_0_ = 0.99 mM glucose) liquid cultures containing varying fosfomycin concentrations show a uniform initial increase in cell biomass followed by growth or death depending on the antibiotic concentration. **C** Similar measurements in nutrient-poor (*c*_0_ = 0.037 mM glucose) cultures shows a slight biomass increase, but no cell death. Fits shown in **B**-**C** directly parameterize the model (Fig. S6, S7). **D-F** Numerical simulations matching the conditions of Fig. **2** recapitulate the dynamics of experimental death fronts; colors are as in Fig. **2**. **G** Representative simulation corresponding to Fig. **1**D recapitulates the progression of the experimental death front, and shows how nutrient consumption sensitizes growing cells (ii), depletes nutrient (iii), and thereby establishes the position of the death front (iv-v). We locate the death front as the furthest position at which 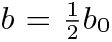, indicated by the vertical dashed lines. All quantities are normalized by their initial values.

As time progresses, both nutrient and antibiotic diffuse through the population with diffusivities *D*_*c*_ and *D*_*a*_, respectively, as described by the first terms on the right hand sides of Eqs. 2-3 (Fig. **3**A). The cells then consume nutrient, with a maximal rate per cell *κ*, following Michaelis–Menten kinetics relative to the characteristic concentration *c*_∗_ independent of antibiotic exposure [62], as described by the second term on the right hand side of Eq. 2. This nutrient consumption leads to growth, with a maximal rate per cell *g*, following Monod kinetics [63–65], as described by the right hand side of Eq. 1a. Growth is also strongly influenced by antibiotic exposure. To quantitatively describe this influence, we directly measure bacterial growth and death across a broad range of glucose and fosfomycin concentrations (Fig. **3**B-C). Our measurements reveal a simple rule: When *a > a*_∗_ ≈ MIC, cells with access to sufficient nutrient grow exponentially to a threshold concentration Φ*b*_0_, with Φ = 1.5, and then die exponentially with a maximal rate *d* (Fig. **3**B)—consistent with previous studies of other antibiotics that also kill bacteria by targeting their cell wall [66]. This effect is quantified by the right hand side of Eq. 1b.

Remarkably, numerical simulations of this minimal model—fully parameterized using separate measurements (Table S1)—show that it recapitulates all the key features of our experimental observations, as summarized by Fig. **3**D-F (compare to Fig. **2**A-C). As in the experiments, when enough antibiotic is present, a death front progressively sweeps through the population, controlled by nutrient transport and availability. The simulations also provide useful information on the underlying cell-scale processes that shape the death front (Fig. **3**G, Movie S6). As nutrient (gold) and antibiotic (purple) diffuse into the bacterial population (green), cells near the surface of the population become metabolically active, consume nutrient, and grow (arrow in panel ii). This process has two key consequences: it causes the nutrient to lag behind the antibiotic (arrow in iii), and it causes the cells to become more sensitive to killing by the antibiotic (dip in the green curve in iii-iv). As a result, a death front (dashed line) forms and progresses through the population (arrows in iv-v), with its motion constrained by the extent to which nutrient can penetrate—the nutrient bottleneck. Ultimately, this sequential process of nutrient transport, cell growth sensitization, killing by antibiotic, and further nutrient transport revealed by the simulations controls death front dynamics.

### Quantitative principles underlying the formation and dynamics of death fronts

The close agreement between our simulations and experiments indicates that that the biophysical picture described in Fig. **3**A captures the essential processes underlying antibiotic killing in structured bacterial populations. Analysis of the model also enables us to establish quantitative principles describing death front dynamics across a broad range of conditions (Table S2). To do so, first, we consider an idealized system in which nutrient depletion is minimal. In this case, the death front forms when both nutrient and antibiotic diffuse into the population and reach the characteristic concentrations *c*_∗_ and *a*_∗_, driving subsequent cell growth sensitization and killing over a characteristic time scale 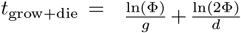. An example analytical calculation quantifying these processes (detailed in *SI Appendix*) to predict death front progression after 12 h is shown by the background color in Fig. **4**A. As expected, when nutrient is more abundant (lower right), death front progression is limited by transport of the antibiotic; conversely, when antibiotic is more abundant (upper left), the death front is nutrient-limited. Our simulations confirm this expectation, as shown by the filled circles.

**Fig. 4.**
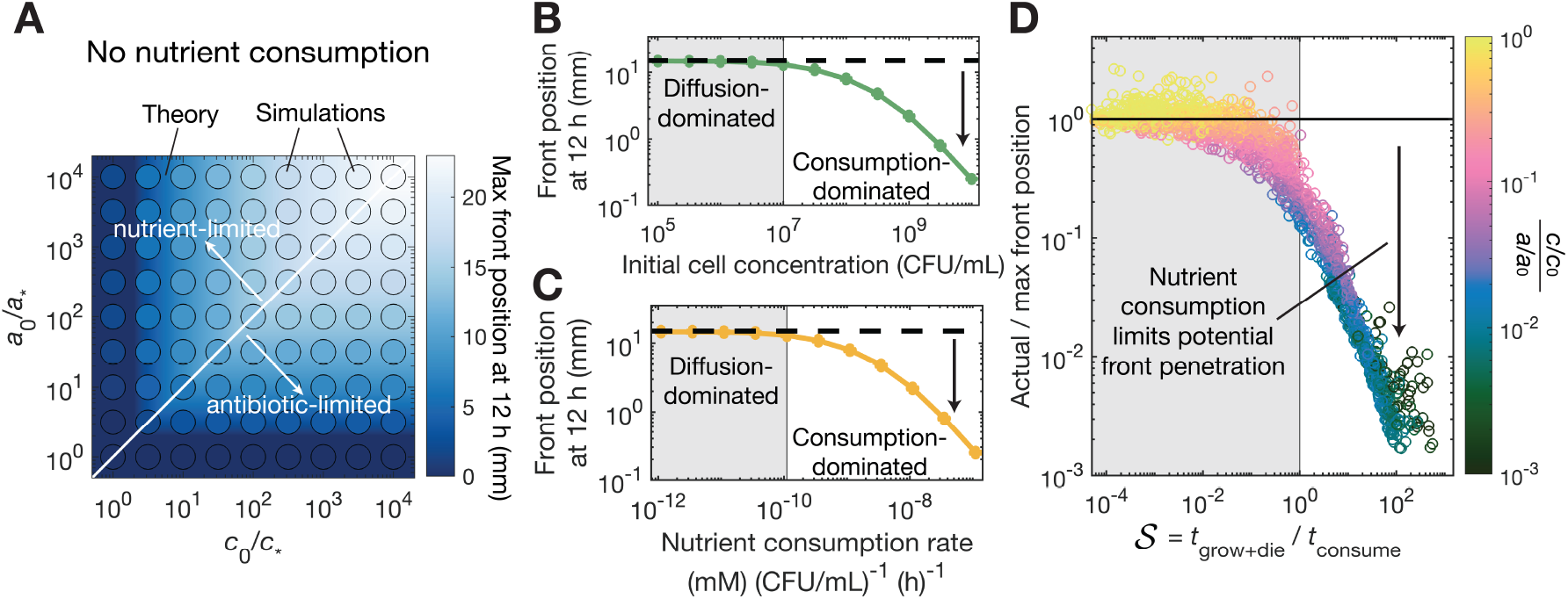
Model reveals when nutrient consumption limits death front propagation. **A** When nutrient depletion is minimal, death front progression is controlled by the diffusive transport of nutrient and antibiotic—as exemplified here by tracking the front position after 12 h for different nutrient (abscissa) and antibiotic (ordinate) concentrations. Background color shows the theoretical prediction (*SI Appendix*) and points show the results of numerical simulation of our model with *κ* = 0. When antibiotic exceeds nutrient (*c*_0_*/c*_∗_ *< a*_0_*/a*_∗_), death front progression is limited by nutrient transport, whereas in the reverse case (*c*_0_*/c*_∗_ *> a*_0_*/a*_∗_), front progression is antibiotic-limited. **B**-**C** Bacterial consumption of nutrient slows death front progression below the theoretical maximum calculated in **A** (dashed black lines). Points show results of numerical simulations with either **B** fixed consumption rate per cell *κ* = 1.1× 10^−9^(mM)(CFU*/*mL)^−1^(h)^−1^ and increasing cell concentration *b*_0_ or **C** fixed *b*_0_ = 10^8^ CFU/mL and increasing *κ*. **D** Data corresponding to **B**-**C** but exploring a vast range of parameter values (Table S2, Fig. S8) all collapse when represented using the slowdown parameter 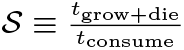. This analysis reveals that when *𝒮 <* 1, the effect of nutrient depletion is minimal, whereas when *S* increases above unity, nutrient depletion (indicated by the color) increasingly hinders death front progression. We choose *D*_*c*_ = *D*_*a*_ and *c*_0_*/c*_∗_ = *a*_0_*/a*_∗_ for this example calculation. Cases where no death front forms after 12 h are not shown.

These predictions represent a theoretical maximum for how far a death front can progress through purely diffusive transport of nutrient and antibiotic (dashed black lines in Fig. **4**B-C). However, bacterial consumption can appreciably deplete nutrient as well, further constraining the progression of the death front, as we found experimentally in Fig. **2**B-C. Our simulations (points in Fig. **4**B-C) recapitulate this effect: as either cell concentration *b*_0_ or the nutrient consumption rate *κ* increase above threshold values (vertical lines), the motion of the death front is increasingly hindered. Therefore, we consider an additional characteristic time scale 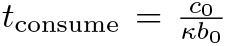, which provides an estimate for how long it takes for the bacteria to deplete nutrients. Comparing *t*_grow+die_ and *t*_consume_ then yields the dimensionless “slowdown” parameter, 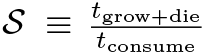. When 𝒮 *<* 1, we expect that the effect of nutrient depletion is minimal, and the death front moves at its maximal rate as given by Fig. **4**A. By contrast, as 𝒮 increases above 1, we expect that nutrient depletion increasingly limits the motion of the death front. To test this prediction, we perform thousands of simulations of our model, exploring a broad range of conditions found in nature (Table S2), and determine the front position after 12 h compared to the diffusion-dominated theoretical maximum. Despite the large variation in conditions, with the underlying parameter values tested ranging across many orders of magnitude, all the data collapse, as shown by the different points in Fig. **4**D. The color indicates the ratio between local nutrient and antibiotic concentrations at the position of the death front; as expected, with increasing 𝒮 *>* 1, nutrient scarcity increasingly constrains the death front. Thus, taken altogether, our work quantitatively describes the conditions under which nutrient consumption by metabolically active cells near the surface of a bacterial population can slow the death front and protect inner cells from antibiotic killing, as has long been conjectured [23–27].

### Lower antibiotic dosage enables heteroresistant regrowth in the wake of a death front

Our analysis thus far has, for simplicity, assumed that the bacterial population is homogeneously sensitive to the antibiotic; that is, the critical level of antibiotic needed to kill cells, *a*_∗_, is the same for the entire population. This assumption is appropriate when antibiotic is administered at a level far above MIC (*a*_0_ ≫ *a*_∗_), and thus, slight cell-to-cell variations in *a*_∗_ have a minimal effect. However, in reality, bacterial populations are often heteroresistant: their antibiotic sensitivity and resistance levels vary across cells, even if they are genetically identical [67–69]. This phemonenon is thought to underlie the failure of many antibiotic treatments—with fosfomycin being a prominent example—in the clinic [70–72]. Indeed, using a standard assay [68], we find that the cells used in our experiments are heteroresistant (Fig. S9): while the characteristic *a*_∗_ for the entire population is ≈ 7 µg/mL, some cells are killed by levels of fosfomycin as low as *a*_∗,min_ ≈ 2 µg/mL, while others survive exposure to fosfomycin concentrations as large as *a*_∗,max_ ≈100 µg/mL. What are the population-scale implications of this heteroresistance?

The approach developed here provides a straightforward way to address this question. In particular, we predict that exposing a structured population to a fosfomycin concentration *a*_0_ between *a*_∗,min_ and *a*_∗,max_ will not achieve complete clearance of the cells. Instead, after the death front sweeps through the population, we expect that subsequent glucose diffusion into the population facilitates regrowth of the resistant cells. Our experiments directly confirm this prediction: as shown for the case of *a*_0_ = 64 µg/mL in Fig. **5**A (Movie S7), a death front initially sweeps through the population (*t <* 12 h), but then, small microcolonies of resistant cells (indicated by the white arrows for *t* ≥ 12 h) regrow in its wake. Testing an even smaller *a*_0_ = 16 µg/mL yields the regrowth of even more resistant microcolonies, additionally confirming our expectation (Fig. **5**B, Movie S8). Finally, exploring the case of *a*_0_ = 64 µg/mL again, but this time with a population that is 10× more concentrated, tightens the nutrient bottleneck and slows the death front—producing more resistant microcolonies in its wake, also as expected (Fig. **5**C, Movie S9). Therefore, while supplying nutrients promotes bacterial killing at large antibiotic dosage, those same nutrients paradoxically promote the selection and regrowth of heteroresistant bacteria, allowing for population recovery, when antibiotic is administered at intermediate levels *a*_∗,min_ *< a*_0_ *< a*_∗,max_. Our visualization also highlights a unique feature of population structure: in well-mixed culture, regrowth by a resistant subpopulation typically occurs by a single cell outcompeting the entire population [73–77], whereas in a structure population, multiple microcolonies can be maintained simultaneously (Fig. **5**), potentially allowing the population to improve its future survival through bet hedging [78–84].

**Fig. 5.**
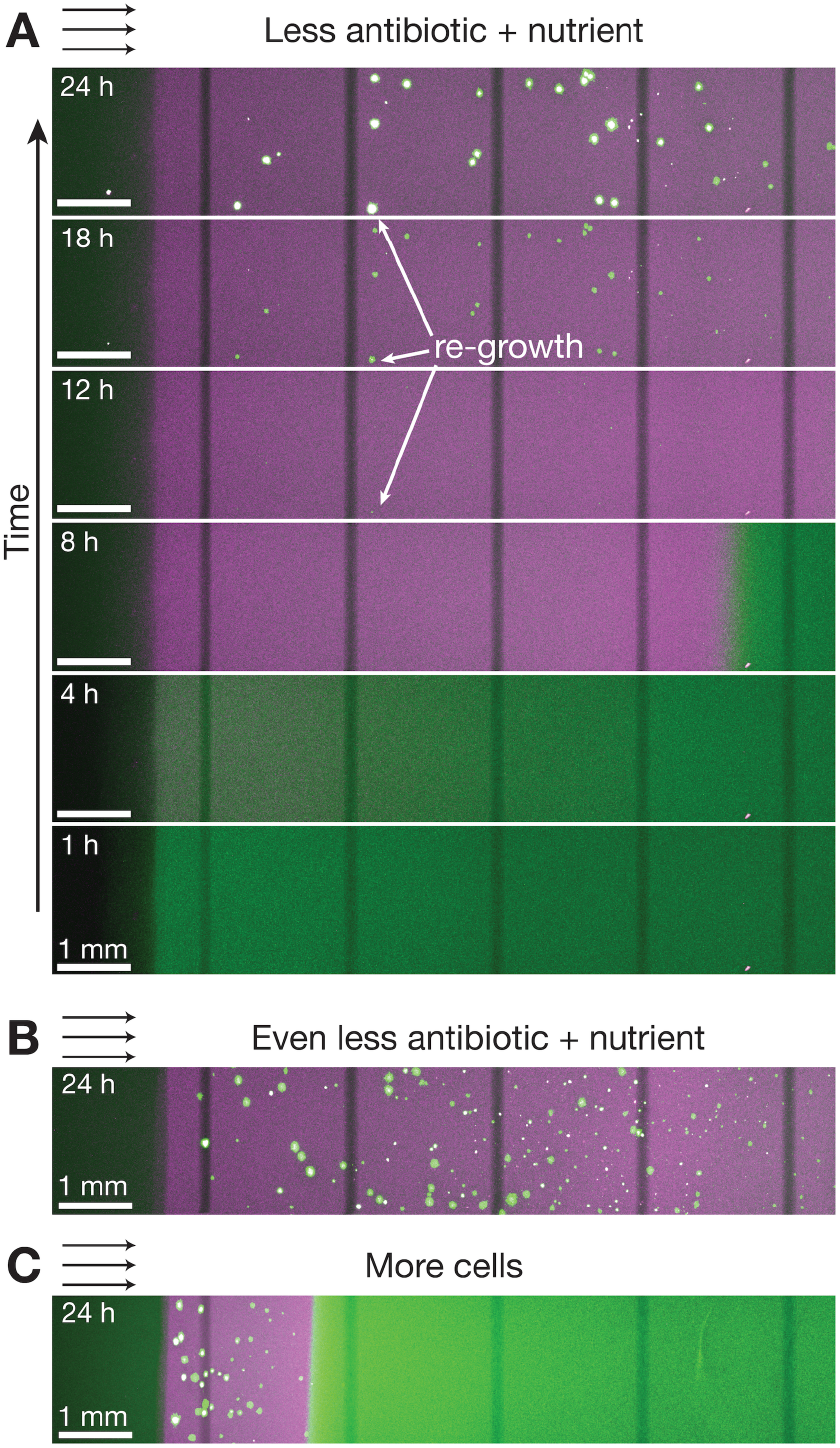
When exposed to lower antibiotic dosage, resistant micro-colonies regrow in the wake of the death front. **A** Same experiment as in Fig. **1**D, but with less fosfomycin (*a*_0_ = 64 µg/mL) and more glucose (*c*_0_ = 2.2 mM). After the death front sweeps through the population (*t <* 12 h), microcolonies of resistant cells regrow in its wake (arrows, *t* ≥ 12 h). **B** Using an even lower fosfomycin concentration (*a*_0_ = 16 µg/mL) leads to more regrowth of resistant microcolonies. **C** Using a higher concentration of cells (*b*_0_ = 10^9^ CFU/mL) also leads to more regrowth of resistant microcolonies, as well as a slower death front. Micrographs show maximum intensity projections of optical slices taken 100 µm apart over the entire mm depth of the sample. Dim vertical stripes are an artifact of stitching multiple imaging fields of view together.

## DISCUSSION

In this study, we have demonstrated that the spatial structure of bacterial populations fundamentally alters their response to antibiotic treatment through the coupling of nutrient and antibiotic transport (at large scales) and cellular metabolism and antibiotic activity (at small scales). Using a well-controlled experimental system with *E. coli* immobilized in transparent hydrogel matrices, we directly visualized the propagation of cell death as a traveling front when exposed to sufficient levels of both glucose (as well as other 6-carbon sugars) and fosfomycin. Our biophysical model (Fig. **3**A) provides a framework to quantitatively describe the occurrence and dynamics of such death fronts across a broad range of cell types, nutrients, and antibiotics (Fig. **4**D); indeed, many different microbial species exhibit similar metabolic-dependent responses to diverse classes of antibiotics [7, 20–22, 85–88]. Our findings thereby expand the typical view of how antibiotics attack natural bacterial populations (Fig. **1**A). In particular, they reveal that nutrient availability serves as a critical bottleneck to antibiotic killing, with collective nutrient consumption by cells at the population periphery transiently creating a protective shield for those in the interior (Fig. **6**A)—as quantified by the condition 𝒮 *>* 1. By contrast, when nutrient consumption is too slow (𝒮 *<* 1), population clearance is limited only by the diffusive transport of nutrient and antibiotic through the population (Fig. **6**B).

**Fig. 6.**
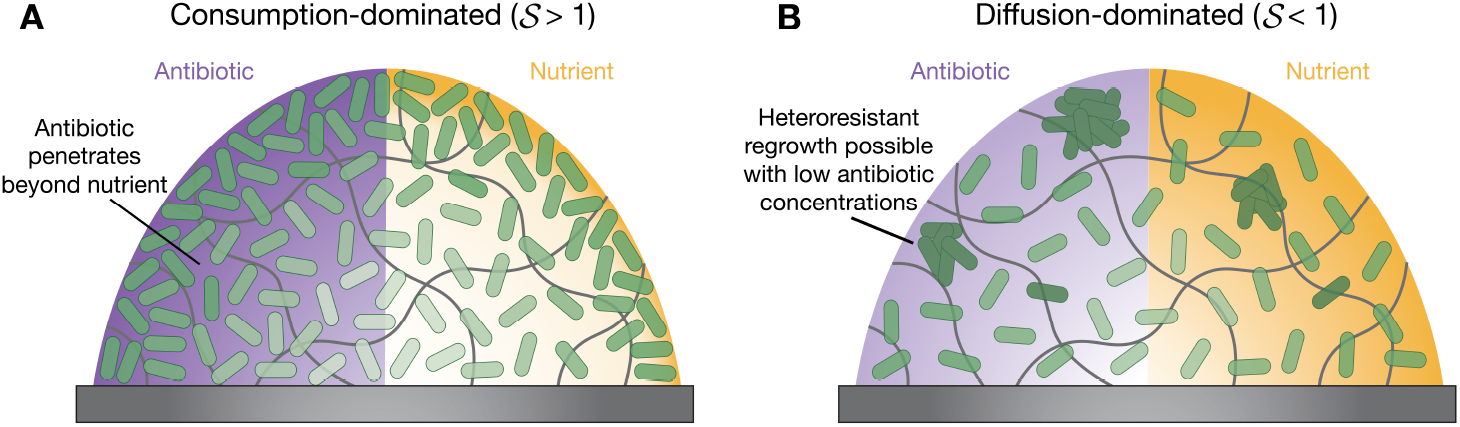
Consumption-dominated vs. diffusion-dominated regimes of antibiotic treatment impact outcomes. **A** When collective nutrient consumption is rapid (*𝒮 >* 1), cells in the interior of the population are nutrient deprived, and thus protected against the antibiotic, even if it can penetrate in. **B** When nutrient consumption is slow (*𝒮 <* 1), death front propagation is limited only by nutrient and antibiotic diffusion.

Our study has several limitations that present opportunities for future research. First, we focused on a single antibiotic (fosfomycin) and cell type (*E. coli*); building on our work to explore other combinations will be a useful way to further probe the generality of our findings. Second, while our hydrogel-based system provides direct optical access and control over population structure and environmental conditions, it does not fully recapitulate the complexity of natural bacterial populations, which often contain multiple species [89, 90] and extracellular polymeric substances [91] that may further modulate nutrient and antibiotic transport. Extending our approach to more complex multispecies communities would help bridge this gap. Third, our current model does not account for nutrient recycling of dead cells [92–94], as well as potential phenotypic adaptations [34] and gene expression changes [95, 96] that may occur during antibiotic exposure, which will be useful to consider in future studies.

Nevertheless, despite these limitations, our findings have broad implications across multiple domains. In clinical microbiology, our work provides a mechanistic basis for the often-observed discrepancy between antibiotic efficacy in liquid culture versus in vivo infections. By understanding how spatial structure and nutrient availability influence bacterial survival, clinicians could design improved treatment protocols that account for these factors—for example, by combining antibiotics with compounds that disrupt population structure or enhance nutrient penetration [97–100]. More broadly, our findings could help predict and control the behavior of structured bacterial populations in the environment (e.g., soil, sediments, and aquatic systems) and industry (e.g., bioremediation and biofuel production).

Our findings also reveal how nutrient-mediated protection extends survival time for cells within structured populations, which has considerable implications for antibiotic resistance. Unlike in well-mixed cultures where only rare persisters survive [101–103], spatial structure enables entire subpopulations to experience extended antibiotic exposure without dying, providing more cells with conditions to adapt and potentially develop resistance, even without net growth [104]. This prolonged survival may also trigger population-wide signaling and phenotypic switching, such as increased biofilm matrix production or enhanced motility and dispersal to new environments [105]. These advantages add to the growing list of other protective mechanisms conferred by population spatial structure that have recently been documented [34, 57, 89, 106– 110]. Building on the fundamental framework established here to investigate these possibilities will be an important direction for future work—not only advancing basic understanding of collective microbial behavior but also offering practical insights for improved antibiotic stewardship in an era plagued by increasing antimicrobial resistance.

## MATERIALS AND METHODS

Cells are grown in either Luria Bertani (LB) broth or M9 media (Difco) supplemented with a single carbon source, namely glucose, glycerol, or mannose. Aqueous fosfomycin stocks are made fresh for each experiment. The hydrogel matrices are prepared by dispersing dry granules of acrylic acid-alkyl acrylate copolymer microgels (Carbomer 980; Lubrizol, Wickliffe, Ohio) in liquid M9 media, mixed for at least 12 h using a rotary mixer, and pH adjusted to 7.0 by adding 10 M NaOH. Immediately prior to each experiment, glucose, glycerol, mannose, fosfomycin, and/or propidium iodide (Sigma Aldrich) are added to each hydrogel matrix as necessary. The hydrogel matrices are then deposited in the wells of a 4 chambered coverglass dish (Cellvis), with a custom-fit acrylic divider placed in the center to separate the cell-containing matrix on one side from the nutrient and antibiotic-containing matrix on the other side. We initiate each experiment by gently removing the divider so the matrices on both sides join together. Finally, we cover the top surface of each sample with *∼* 0.5 mL paraffin oil to prevent evaporation during imaging. The bacterial populations are imaged every 30− 60 min using a Nikon A1R+ inverted laser scanning confocal microscope with a temperature-controlled stage maintained at 37°C. Multiple images are stitched together to image the entire cell population with a resolution of ≈ 900 nm/pixel. For the micrographs in Fig. **5**, we image at a resolution of 2.2 µm/pixel and take *∼* 35 optical slice images spaced 100 µm apart.

## Acknowledgment

We acknowledge support from National Science Foundation (NSF) grants CBET-1941716, DMR-2011750, and EF-2124863 as well as the Camille Dreyfus Teacher-Scholar and Pew Biomedical Scholars Programs, the Eric and Wendy Schmidt Transformative Technology Fund, and the Princeton Catalysis Initiative. This work was supported in part by the NSF Graduate Research Fellowship Program (to A.M.H.) under grant no. DGE-2039656; any opinions, findings, and conclusions or recommendations expressed in this material are those of the authors and do not necessarily reflect the views of the NSF. We thank Katherine Sniezek for constructing the strain used in these experiments and for instructions on the population analysis profiling assay, as well as Mark Brynildsen, Bruce Levin, Ned Wingreen, the late Kevin Wood, Lingchong You, and members of the Datta Lab for stimulating discussions and useful feedback.

## Supporting information (SI)

### Experimental details

#### Bacterial strains, growth media, and antibiotics

All experiments are conducted using *E. coli* K-12 substrain MG1655 ΔlacI ΔaraBAD :: P_T5_ − gfp− kanR, which constitutively expresses GFP from the chromosome and whose construction has been described previously [1]. Cells are grown in either Luria Bertani (LB) broth or M9 media supplemented with a single carbon source, namely glucose, glycerol, or mannose. LB broth is made by dissolving 2% (w/v) LB powder (Sigma Aldrich) in Milli-Q water for a final concentration of 0.5 g/L sodium chloride (NaCl), 10 g/L tryptone, 5 g/L yeast extract and sterilized by autoclaving. M9 Media is made by dissolving M9 salts (Difco) in MilliQ water and sterilized by autoclaving. After autoclaving, filter-sterilized magnesium sulphate is added such that the final salt concentrations are 33.9 g/L disodium phosphate (anhydrous) (Na_2_HPO_4_), 15 g/L monopotassium phosphate (KH_2_PO_4_), 2.5 g/L sodium chloride (NaCl), 5 g/L ammonium chloride (NH_4_Cl), and 0.241 g/L magnesium sulphate (MgSO_4_). Immediately prior to each experiment, filter-sterilized glucose, glycerol, or mannose is added to the reported concentration as a carbon source. Fosfomycin stocks are made fresh for each experiment by dissolving fosfomycin-disodium salt into autoclaved Milli-Q water and filter-sterilizing. Then, stocks are diluted appropriately into either liquid media or pre-swollen granular hydrogels depending on the experiment.

#### Preparing the granular hydrogel matrices

We use dense packings of hydrogel grains (“microgels”) as growth matrices for bacteria. Each matrix is prepared by dispersing dry granules of internally cross-linked microgels made of biocompatible acrylic acid-alkyl acrylate copolymers (Carbomer 980; Lubrizol, Wickliffe, Ohio) in liquid M9 media (without carbon sources or antibiotics). The granules absorb the liquid until their elasticity prevents further swelling. We ensure a homogeneous dispersion of swollen microgels by mixing for at least 12 h using a rotary mixer, and adjust the final pH to 7.0 by adding 10 M NaOH. Immediately prior to the experiments, glucose, glycerol, mannose, or fosfomycin disodium are added to the granular hydrogel at reported concentrations. The swollen microgels are ∼5-10 µm in diameter with ∼ 20% polydispersity, but have an internal mesh size of ∼ 40–100 nm, which permits small molecules such as fosfomycin, glucose, and oxygen to freely diffuse throughout while impeding cellular motion [2]. Thus, cells are trapped in the pores of the granular hydrogel matrix to either grow or die in place, but eliminating any effects of chemotaxis or cell motility.

#### Creating model structured bacterial population

We construct tunable and well-defined structured bacterial populations by dispersing stationary phase *E. coli* within granular hydrogel matrices and patterning them next to a reservoir of nutrients and antibiotics in the same granular hydrogel matrix. First, we grow cells to stationary phase to be resuspended into the swollen granular hydrogel media. Two days prior to each experiment, we pick a single *E. coli* colony from an agar plate and grow it in 2 mL of LB for 24 h in a 37°C shaking incubator. After 24 h, we use 200 µL of this 2 mL culture to inoculate 20 mL of LB, and this larger cell culture is also grown for 24 h in a 37°C shaking incubator, establishing a large volume of stationary phase cells which are used in the experiments. After 24 h of growth, we measure the optical density the stationary phase culture, and use this value to determine the exact volume of cells to spin down for the experiment to reach our desired final cell concentration, since an equivalent optical density of 0.15 corresponds to 10^8^ CFU/mL and an equivalent optical density of 1.5 corresponds to 10^9^ CFU/mL. We spin the appropriate volume of cells, which ranges from 100 µL - 4 mL depending on the experimental condition, for 15 minutes at 3000 RCF and resuspend the pellet in ∼ 30 µL of the supernatant. Then, we add this entire cell slurry into an aliquot of the granular hydrogel matrix and evenly disperse the cells by manual shaking to reach the final desired cell concentration. We remove the bubbles introduced by mixing by centrifuging at 3000 RCF for 10 s. This evenly dispersed cell mixture constitutes our model structured cell community. To prepare the nutrient and antibiotic reservoir, we add sterile glucose and fosfoymcin stocks into M9 granular hydrogel matrix to their final concentrations for the reservoir. We also add propidium iodide (Sigma Aldrich) to a final concentration of 6 mM in the cell and antibiotic reservoir to act as a dead cell signal.

To pattern the cell community and reservoir, we place a custom fit, 3 mm acrylic divider in the center of each well of a 4 chambered coverglass dish (Cellvis). We gently deposit 0.4 mL of the cell mixture on one side of the divider and 0.4 mL of the nutrient antibiotic reservoir on the other side of the divider with 1 mL syringes with 20 gauge needles attached. Then, we remove the acrylic divider so the two sides collapse together create a smooth interface over which the antibiotics and nutrients can diffuse into the cell population, marking *t* = 0 h for the experiment. Finally, we cover the top surface of the gel with with *∼*0.5 mL of parrafin oil to prevent evaporation during imaging.

We confirm the initial cell concentration for each experiment by measuring the colony forming units of the cell mixture within the granular hydrogels. We remove ∼ 100 µL of granular hydrogel and dilute it into 500 mL of PBS in a 1.5 mL tube. We confirm this initial dilution volume by massing the tube before and after adding granular hydrogel. Then, we perform serial 10X dilutions in phosphate buffered saline (PBS) before plating 3, 10 µL droplets of each dilution onto a plate containing 1.5% agar, 2% LB. We incubate the plates at 30°C for ∼16 h before counting colonies and converting to CFU/mL.

#### Imaging structured bacterial populations and tracking death front position

All experiments are imaged every 30-60 minutes using a Nikon A1R+ inverted laser-scanning confocal microscope maintained at 37°C. Multiple images are stitched together to image the entire cell population with a resolution of 0.9 µm/pixel. To image the death front progression, 3 separate z slices are imaged at 50, 100, and 150 µm above the bottom surface of the population and then merged. The death front position is defined as the position at which dead cell propidium iodide signal exceeds a threshold value that is held constant across experimental nutrient and antibiotic conditions for the same cell density. When a death front is not reported, no spatial location ever exceeds the threshold signal value. For imaging heteroresistance, we image at a resolution of 2 µm/pixel and take ∼ 35 z slice images spaced 100 µm apart to capture the entire 3D population over time.

#### Measuring well-mixed growth and death rates to parameterize the model

To understand the joint effects of nutrients and antibiotics on cell growth and death, we perform a series of well-mixed experiments using a Biotek Epoch 2 microplate spectrophotometer. First, cells are grown to stationary phase following the same protocol as for creating the structured bacterial population. Then, we wash the stationary phase cells twice with M9 media by centrifuging for 15 minutes at 3000 RCF, removing the supernatant, and rususpending the pellet into M9 media, to remove any residual nutrients from the overnight growth. After the second wash, we further dilute the washed cells with M9 media to 10X the initial desired concentration for the experiment. We prepare a 96 well plate with serial dilutions of M9 media with defined concentrations of glucose and/or fosfomycin. Then, we inoculate these wells with 15 µL of washed cells for a final well volume of 150 µL. We characterize the bacterial growth and/or death dynamics by incubating the microplate at 37°C with continuous linear shaking at 567 cycles per min over a distance of 3 mm for 16-48 h, measuring UV-vis absorption at 600 nm (OD_600_) every 10 min. Finally, we verify the starting concentration of the washed cell inoculum by serially diluting and plating cells on 2% LB 1.5% agar before incubating overnight and counting colonies, similar to the protocol for experiments on structured bacterial populations.

#### Testing for antibiotic degradation

To test if there is any loss of potency for the antibiotic from incubation at 37°C in M9 media, we repeat bacterial growth and death measurements in fosfomycin that was pre-incubated compared to fresh stock (Fig. S3). Both 48 and 24 hours prior to the experiment, we prepare a stock solution of fosfomycin, filter-sterilize it, and dilute it to our working concentration in M9 media supplemented with 24.4 mM glucose (which would dilute to 22.2 mM glucose after cell stocks are added). We incubate the fosfomycin-glucose M9 media at 37°C in a stationary incubator for 48 and 24 h respectively. Then, we prepare serial dilutions with the pre-incubated stocks as well as freshly prepared fosfomycin in a 96-well microplate. We inoculate these cultures with washed stationary phased cells of two different densities which are prepared as previously described for other microplate measurements. Following inoculation, we incubate the microplate at 37°C with continuous linear shaking at 567 cycles per min over a distance of 3 mm for 16-48 h, measuring UV-vis absorption at 600 nm (OD_600_) every 10 min. We compare the growth and death dynamics for cells in freshly made or old fosfoymcin and saw no difference, indicating no degradation or loss of potency over the time scales of our experiment.

#### Quantifying heteroresistance

The heteroresistance profile of the cells, or the resistant subpopulations able to grow at varying antibiotic concentrations, was quantified using standard methods of population analysis profiling [3, 4]. Briefly, cells are grown to stationary phase following the same protocol as for creating the structured bacterial population or measuring well mixed growth and washed once with M9 media. Then, we perform 10X serial dilutions of the washed cells and plate 3, 10 µL droplets each 10X dilution onto plates containing 1.5% agar, M9 media with 22 mM glucose, and 2 fold increments of fosfomycin ranging from 0.5-512 µg/mL. The number of resistant mutants is compared to cells from the same culture grown on antibiotic-free plates to give the frequency of resistant bacterial subpopulations. Resistant subpopulations with frequencies greater than 10^−7^ were detected over an 8-fold concentration range of fosfomycin, marking this cell strain heteroresistant to fosfomycin.

#### Developing, parameterizing, and simulating the model

We use the experiments described below to formulate and parameterize the following 1D continuum model of our experiments.

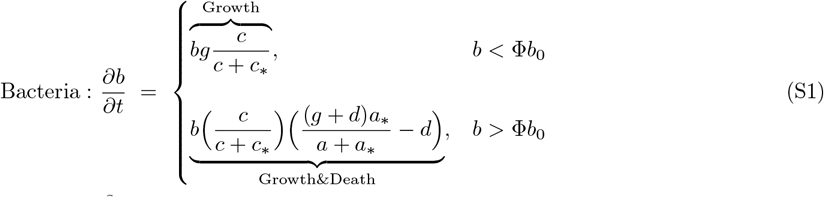

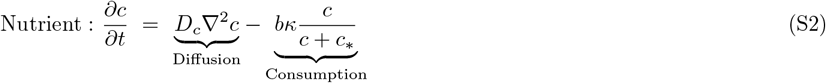

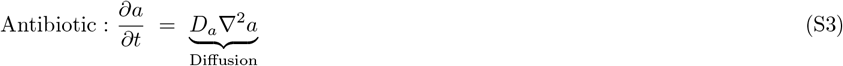

Since the interstitial mesh of our granular hydrogel (∼100 nm) is much larger than both glucose and fosfomycin (∼1 nm), we approximate the diffusivity of each of these small molecules to be uniform and equivalent to the diffusivity of glucose in water *D*_*c*_ = *D*_*a*_ [5], as has been done previously for granular hydrogel systems [6–8]. For the antibiotic, there is no substantial degradation or loss of potency over the timescale of our experiments (Fig. S3), and the effects of cell uptake are negligible (see calculation below), so the dynamics of the antibiotic are entirely captured by diffusion (Eq. 3). In contrast, nutrients are depleted by cells in the system. The growth of bacteria on single carbon sources is well understood by Monod kinetics [9–11], and this growth and consumption interaction forms the basis for our model, as detailed further below.

#### Maximum growth rate

We measure the maximum exponential growth rate of *E. coli* grown in M9 media with 22 mM glucose, as shown in Fig. S4. We fit an exponential curve to the region over hours 4-9 for 8 replicates across four independent experiments to obtain an average maximum growth rate of 0.32 h^−1^.

#### Nutrient consumption rate per cell

Next, we fit the nutrient consumption rate per cell using yield measurements. We grow cells cultures with varying initial cell density in a range of glucose concentrations, following the protocol for plate reader growth. Then, after 20 h of growth in a shaking plate reader, we measure the cell density (CFU/mL) of each sample, as shown in Fig. S5A. To estimate the nutrient consumption rate per cell from these measurements, we again turn to the well established Monod model for bacterial growth on single, consumable carbon sources. Bacteria *b*(*t*) grow from an initial density *b*_0_ according to the equation 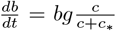, where *g* is the maximum growth rate and *c*(*t*) is the local nutrient concentration. Nutrients *c*(*t*) are simultaneously depleted from their initial concentration *c*_0_ according to equation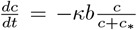, where *κ* is the maximum consumption rate per cell. For simplicity, we assume that cells are always growing and consuming nutrients at the maximal rates *g* and *κ*, thus setting the 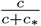 Monod term to 1. Thus, bacterial density over time obeys the equation 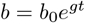. Cell reach a final density of *b*_*f*_ in time 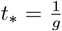ln 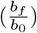. Assuming that cells reach this final density once they have completely exhausted nutrients from initial value of *c*_0_ to 0, we can integrate the equation for nutrients 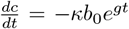 from *c*_0_ to 0 in the time 0 to *t*_∗_. By solving for *κ*, we obtain that 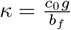. Calculating *κ* from these measurements in Fig. S5A gives a value of *κ* = 2.0 *×* 10^−11^ *±* 2.3 *×* 10^−12^ (mM)(CFU/mL)^−1^(h)^−1^.

To confirm that this value matches the dynamics we see in plate reader growth, we compare simulated well mixed growth to independent experiments measuring cell growth over time in a range of glucose concentrations. First, we convert the OD_600_ values from the plate reader to CFU/mL using a linear conversion rate (Fig. S5B). Then, we simulate well-mixed growth according to unapproximated Monod kinetics across 4 nutrient conditions (Eqn. 1-3 without diffusion terms) and compare these results to experimental measurements seeing good agreement with our cell yield estimation.

In the model, we assume that the consumption rate of nutrients is not impacted by local antibiotic concentration, based on prior work [12]. For simplicity, we also omit any lag time and assume that cells begin growing and consuming nutrients as soon as they encounter glucose levels *c > c*_∗_.

#### Growth threshold

To understand how the presence of fosfomycin influences cell growth and death rates across nutrient conditions, we culture bacteria in a range of glucose and fosfomycin concentrations and measure cell growth and death via UV vis spectroscopy. In high nutrient environments, as shown in Main Text Fig. 3B, we observe that at first cells in both antibiotic rich and poor environments grow exponentially (*t* = 0 − 2 h). Only after growing to a threshold cell density Φ*b*_0_ do cells begin to die exponentially in high antibiotic environments (*a > a*_∗_), similar to what has been previously reported for *β*-lactam antibiotics which also target the cell wall [13]. If lower but non-zero initial nutrients in the culture cannot support a biomass increase to Φ*b*_0_, no death is observed, even in high antibiotic environments (Main Text Fig. 3C). Thus, the first rule for bacteria in our model is that, initially, cells grow exponentially according to only Monod kinetics until reaching the threshold density (Eq. 1a). The growth threshold parameter Φ thus describes the threshold cell density (relative to the initial cell density) that cells must grow to in order to “feel” the effects of local antibiotic and transition from an initial period of exponential growth to a period of exponential death.

To determine the value of Φ, we culture cells in a range of nutrient concentrations for 0 and 256 µg/mL Fos-Na. We compare initial cell density to final cell density and identify the critical initial nutrient level *c*_crit_ below which no cell death is observed for each initial cell density (dashed lines in Fig. S6A, y-axis values of dots in Fig. S6B). As expected, the critical nutrient level is linearly dependent on initial cell concentration *b*_0_ (Fig. S6B) and we can use this linear relation between *b*_0_ and *c*_crit_ to parameterize Φ according to the following calculations.

According to the well-established model of Monod kinetics used above to calculate nutrient consumption rate per cell, we can calculate the time *t*_crit_ that it takes for bacteria to grow from an an initial density *b*_0_ to a final density Φ*b*_0_. Assuming that bacteria always grow at the maximal exponential rate 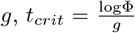. During this period of growth, bacteria are likewise depleting nutrients according to the equation 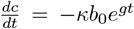. The minimum initial nutrient concentration *c*_crit_ that supports the growth of the population from *b*_0_ to Φ*b*_0_ is the amount of nutrients depleted by the growing population in time *t*_crit_. Thus, we can integrate this equation from time *t* = 0 to *t* = *t*_crit_ where nutrients drop from an initial concentration of *c* = *c*_crit_ to *c* = 0 and solve for *c*_crit_ to obtain the expression 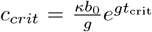. By substituting for 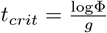, we obtain the linear expression that 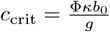. Computing Φ for each experiment (1 dot in Fig. S6B represents 1 experiment shown in Fig. S6A), gives Φ = 1.5 *±* 0.3.

Finally, we simulate well-mixed growth (Eq. 1-3 from the main text, omitting diffusion) using the measured value of Φ = 1.5 for two different cell densities and compare these simulation results to experimental time-course measurements, seeing good qualitative agreement (Fig. S6C). In the simulations, cells reach the threshold density sooner than in experiments, which is expected because our simulations do not include a lag time during which cells grow below their maximum exponential growth rate.

#### Critical antibiotic concentration and maximum death rate

To understand the joint effects of local nutrient and antibiotic concentration on cell growth and death, we culture cells in a range of nutrient and antibiotic concentrations and measure optical density with UV/Vis spectroscopy. In all conditions where initial glucose *c*_0_ is larger than *c*_crit_, we first observe a uniform biomass increase no matter the local antibiotic concentration (Main Text Fig. 3A and Fig. S6C). After this initial period of growth, cells either continue growing exponentially at a maximal growth rate *g* for low antibiotic concentrations or begin dying exponentially at a maximal death rate *d* for high antibiotic concentrations. We define the net exponential growth or death rate as *γ* where 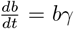. In the limit of no antibiotics, growth is only dependent on nutrient concentration, so 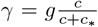, or traditional Monod kinetics.

To measure *γ* = *f* (*c, a*) across experimental conditions, we fit an exponential curve over sliding 2 hour time windows between 1.5 and 9 h of growth for each experimental replicate in each nutrient-antibiotic condition. If any of the measured rates over the sliding time window are less than −0.1h^−1^, we consider that cells die exponentially and take the lowest measured rate over the time window as the reported net growth and death rate *γ*. Next, if biomass does not change, or all measured rates fall between − 0.1 −0.1h^−1^, we take the average rate over the time window to be *γ*. We choose these thresholds to account for the noise that occurs when measuring optical density of dilute bacterial suspensions. Finally, if there is a positive biomass increase over this time window, we report the maximum rate as *γ* for that condition. All measured net growth and death rates *γ* across conditions can be seen in the dots in Fig. S7, which represent 192 replicates across over 96 glucose and fosofmycin conditions for each cell density.

Upon inspection of the data, we first observe that below a threshold nutrient concentration no growth or death is observed (Fig. S7, white points below horizontal dashed line). Above this threshold nutrient concentration, we measure a positive *γ* (growth) at low antibiotic concentrations (Fig. S7, green points) and a negative *γ* (death) at high antibiotic concentrations (Fig. S7, purple points). The threshold nutrient concentration corresponds to *c*_0_ = *c*_crit_ that supports growth of *b*_0_ to Φ*b*_0_ for each cell density we tested (Fig. S7A vs. S7B). The transition between exponential growth and death occurrs at a consistent critical antibiotic concentration *a*_∗_ across nutrient levels (Fig. S7, vertical dashed line). To describe this transition from exponential growth to death in the limit of high nutrient levels 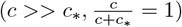, we use an inverse Monod function for antibiotic namely that 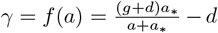. Combining the nutrient and antibiotic dependence for *γ* by multiplying the two Monod terms yields 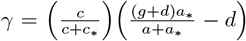 and 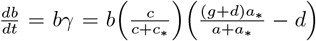 which effectively describes the net bacterial growth or death rate in terms of local nutrient and antibiotic concentration. The multiplication of these two Monod functions follows prior literature describing the joint effects of nutrient and antibiotic [14] and incorporates the broader microbiological phenomenon that maximum death rate varies linearly with maximum growth rate [15].

To parameterize *a*_∗_ and *d* from these data, we fit the data for each cell density to the term 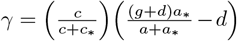 (Fig. S7, background color). We fix the maximum growth rate *g* = 0.32 h^−1^ and obtain fit values for *a*_∗_, *d*, and *c*_∗_. For the simulations, we use parameters from the low cell density data set because they more accurately measure faster maximum growth and death rates at extreme conditions whereas the high cell density data is bounded by the saturating values in optical density measurements. Notably, however, *a*_∗_ does not change appreciably across cell density. Additionally, since the acquired data actually fits *c*_crit_ not *c*_∗_ for the critical nutrient value (increases ∼10 fold for a 10 fold cell density change), we use the well established literature Monod value for *E. coli* growth on glucose *c*_∗_ = 0.02 mM in our simulations [9]. Indeed, UV-vis measurements are not sensitive enough to detect a biomass increase for initial cell populations dilute enough to appreciably grow on *c*_0_ = *c*_∗_. A summary of all parameter values that match our experimental conditions can be found in Table S1.

#### Estimating antibiotic loss into cells

We estimate the cell concentration dependent loss of fosfomycin from the bulk as follows. First, prior work demonstrates that intracellular fosfomycin concentration equilibrates to the extracellular concentration within minutes of antibiotic exposure [16]. Thus, the fraction of intracellular fosfomycin compared to the bulk concentration can be approximated by the volume fraction occupied by cells. A single *E. coli* cell has a volume of approximately 1 µm^3^, so a dense cell population of 10^9^ CFU/mL, 10X denser than the cell concentrations tested experimentally, occupies a total volume fraction of just 0.1%. Thus, only 0.1% of applied fosfomycin is lost into cell cytoplasm by this estimation. Additionally, if we consider that fosfomycin binds irreversibly to the intracellular target protein, MurA [17], we can approximate the molecules of fosfoymin lost into cells as the total concentration of MurA proteins. In exponential phase, cells express the maximum concentration of *∼* 1000 MurA/cell [18]. So, a 10^9^ CFU/mL cell population has *∼* 10^12^ molecules MurA/mL which equates to 0.0002 µg/mL bound fosfomycin compared to bulk concentrations of 16-2000 µg/mL fosfomycin in our experiments. By both of these metrics, the concentration of fosfomycin lost into cells, even at high cell densities, is negligible. Thus, we omit cell uptake terms from our model and entirely capture the dynamics of fosfomycin by diffusion.

#### Numerically simulating the model

To simulate the continuum reaction-diffusion equations of our system, we discretize our system and implement a forward-time, centered-space scheme of finite differences. For all simulations, the temporal and spatial resolution of the simulations is 1 × 10^−4^ h and 0.025 mm, respectively. Repeating representative simulations with varying spatial and temporal resolution reveals that finer discretization does not appreciably alter the results (Fig. S10). Thus, our choice of discretization is sufficiently resolved such that the results are not appreciably influenced by discretization.

For numerical simulations exactly recapitulating the experiments (Main Text Fig. 3E-G), we set the total simulation domain size *L* to 19.9 mm, matching the dimension of the experimental dish and run simulations for 24 h, matching the duration of imaging. For simulations sweeping across biophysical conditions (Main Text Fig. 4A-D), we use a larger system size *L* of 80 mm, ensuring that *L/*2, the size of the bacterial population and maximum position of the death front, is always larger than the simulated death front position at the final time points so that edge effects are minimal. Further, when evaluating front position 12 h after forming, we vary the total simulation time to be either 24 h or at least 20 h after a predicted death front would appear (see below for calculations of the predicted time for a front to appear) for cases with slower growth and death rates. Finally, to match the closed nature of our experimental system, we impose no flux boundary conditions on each edge of the simulated dish.

When sweeping over many biophysical conditions, we randomly sample parameter values across the possible range shown in Table S2, while applying the following constraints. First we restrict that cells must die faster than they grow (*d < g*) to prevent biomass accumulation. Next, *c*_0_*/c*_∗_ and *a*_0_*/a*_∗_ must greater than or equal to 5 to ensure possible theoretical front formation. While non-linearity in the Monod function allows for modest death front formation at lower *c*_0_*/c*_∗_ and *a*_0_*/a*_∗_ values, the theoretical maximum front prediction breaks down. Finally, for data shown in Main Text Fig. 4D, *D*_*c*_ = *D*_*a*_ and *c*_0_*/c*_∗_ = *a*_0_*/a*_∗_ such that the theoretical maximum front position is neither antibiotic nor nutrient limited, rather falling along the white diagonal line of Fig. 4A. When freely simulating all parameters without these constraints, the data still collapse according to our predicted 𝒮 parameter, though less tightly (Fig. S8). In all cases, we define the death front position to be the deepest spatial location where 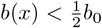 at each time point. All numerical simulations were performed using MATLAB R2024a.

#### Theoretical calculation of death front dynamics

To predict how far a death front could theoretically penetrate into a structured bacterial population, we considered how far both nutrients and antibiotics can diffuse from the reservoir into the cell population in a given amount of time. The concentration profile for a solute diffusing between 2 semi-infinite planar domains with concentrations *c*_1_ and *c*_2_ (*c*_1_ *> c*_2_) can be described by the following function: 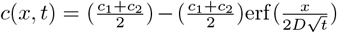 where 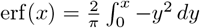 *dy*. For the experimental system geometry, *c*_1_ = *c*_0_ and *c*_2_ = 0. Thus the diffusive profile of nutrients and antibiotics over the system geometry −*L/*2 *< x < L/*2 can be described as: 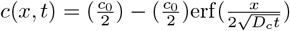.

Without nutrient consumption or other loss, cells will start the growth sensitization and death process at a given position once both nutrients and antibiotics have reached that position locally. Since both cell growth and death obey Monod functions, which sigmoidally step up from a value of 0 to 1 at the critical concentration *c*_∗_, we approximate that growth sensitization and death occur once the local concentration reaches *c*_∗_. A death front is detected at a given position once cells at that position have grown exponentially from an initial concentration *b*_0_ to the growth threshold *ϕb*_0_ and then died exponentially from *ϕb*_0_ to a local cell density of 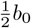, at which point a death front is detected. The total time delay for growth and death *t*_grow+die_ can be approximated as the characteristic time for growth *t*_*g*_ plus the characteristic time for death *t*_*d*_. For simplicity, we assume that cells always grow and die at their maximum exponential rates. Then, exponentially growing cells obey the function 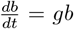. To find *t*_*g*_, we integrate each side of the equation 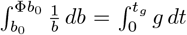 and solve for 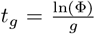. Similarly, exponentially dying cells obey the function 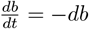. To find *t*_*d*_, we integrate each side of the equation 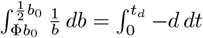 and solve for 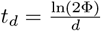. Thus, 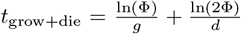. Then to find the maximum death front position after a given time *T*, we solve for the position *x*_*front*_ where *c*(*x*_front_, *t*_eval_) = *c*_∗_ where *t*_eval_ = *T*− *t*_grow+die_ to account for the time delay of growth and death. Since both nutrient and antibiotic must reach their local threshold concentrations for the death front to reach a given position, we take the minimum diffusion distance between the two solutes as the maximum possible death front *x*_front_ position after a given time *T*.

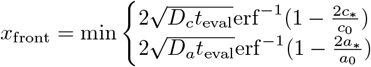

As shown in Main Text Fig. 4A, we see good agreement between this theoretical prediction and numerical simulations of our model. The nonlinearity of the Monod function in the simulation means a front can form and progress even if *a*_0_ *<* 2*a*_∗_ or *c*_0_ *<* 2*c*_∗_, and thus the theoretical predictions slightly deviate for low 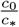 and 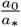.

#### Deriving the dimensionless slowdown parameter 𝒮

To understand exactly how and when nutrient consumption impacts death front propagation, we consider two characteristic time scales: the time for the initial cell population to deplete source nutrients *t*_consume_ and the time scale of growth sensitization and death *t*_grow+die_. We compare these two timescales to obtain a dimensionless death slowdown parameter 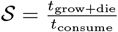.

We previously calculate the term 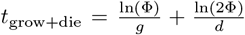 in the section above. Now, to calculate *t*_consume_, we consider the time it takes for a fixed cell population to deplete local nutrients. Bacteria *b*(*t*) grow from an an initial density *b*_0_ according to the equation 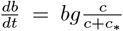. Nutrients *c*(*t*) are simultaneously depleted from their initial concentration *c*_0_ according to equation 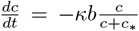. For simplicity, we assume that the cell population size is fixed at *b* = *b*_0_ and that these cells are always are always consuming nutrients at the maximal rates *κ*, thus setting the 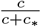 Monod function to 1. So, the nutrient concentration over time obeys the equation 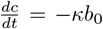. Next, we integrate each side to find the time *t*_consume_ that it takes to deplete nutrients from an initial concentration *c*_0_ to 0: 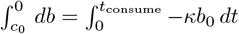. Solving for *t* gives 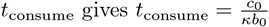.

The dimensionless slowdown parameter 𝒮 compares these two timescales such that 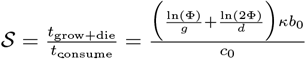 When 𝒮 is small, cells grow and die must faster than they consume and deplete nutrients so the death front can progress at its maximal diffusive pace. When 𝒮 is large, nutrients are consumed and depleted faster than cells can grow and die. Thus, cells are locally starved and can remain tolerant despite high local antibiotic concentration, effectively slowing down bacterial clearance by the antibiotic—the nutrient bottleneck effect.

#### Supplementary movies

Supplementary movies are available on Zenodo (10.5281/zenodo.14990205).

**Movie S1**. Time-lapse microscopy of a 10^8^ CFU/mL cell population being treated with 2048 µg/mL Fos-Na and 0 mM glucose. Green GFP signal corresponds to live cells with intact cell membranes. Magenta signal is propidium iodide, a dead cell indicator. Three biological replicates are shown and scale bar is 1 mm.

**Movie S2**. Time-lapse microscopy of a 10^8^ CFU/mL cell population being treated with 2048 µg/mL Fos-Na and 0.22 mM glucose. Green GFP signal corresponds to live cells with intact cell membranes. Magenta signal is propidium iodide, a dead cell indicator. Three biological replicates are shown and scale bar is 1 mm.

**Movie S3**. Time-lapse microscopy of a 10^8^ CFU/mL cell population being treated with 256 µg/mL Fos-Na and 0.22 mM glucose. Green GFP signal corresponds to live cells with intact cell membranes. Magenta signal is propidium iodide, a dead cell indicator. Three biological replicates are shown and scale bar is 1 mm.

**Movie S4**. Time-lapse microscopy of a 10^8^ CFU/mL cell population being treated with 2048 µg/mL Fos-Na and 2.2 mM glucose. Green GFP signal corresponds to live cells with intact cell membranes. Magenta signal is propidium iodide, a dead cell indicator. Three biological replicates are shown and scale bar is 1 mm.

**Movie S5**. Time-lapse microscopy of a 10^9^ CFU/mL cell population being treated with 2048 µg/mL Fos-Na and 0.22 mM glucose. Green GFP signal corresponds to live cells with intact cell membranes. Magenta signal is propidium iodide, a dead cell indicator. Three biological replicates are shown and scale bar is 1 mm.

**Movie S6**. Numerical simulation of a 10^8^ CFU/mL cell population being treated with 2048 µg/mL Fos-Na and 0.22 mM glucose. Y axis is normalized by initial concentration of cells, nutrient, and antibiotic.

**Movie S7**. Time-lapse microscopy of a 110^8^ CFU/mL cell population being treated with 64 µg/mL Fos-Na and mM glucose. Green GFP signal corresponds to live cells with intact cell membranes. Magenta signal is propidium iodide, a dead cell indicator. Three biological replicates are shown and scale bar is 1 mm.

**Movie S8**. Time-lapse microscopy of a 10^8^ CFU/mL cell population being treated with 16 µg/mL Fos-Na and 2.2 mM glucose. Green GFP signal corresponds to live cells with intact cell membranes. Magenta signal is propidium iodide, a dead cell indicator. Three biological replicates are shown and scale bar is 1 mm.

**Movie S9**. Time-lapse microscopy of a 10^9^ CFU/mL cell population being treated with 64 µg/mL Fos-Na and 2.2 mM glucose. Green GFP signal corresponds to live cells with intact cell membranes. Magenta signal is propidium iodide, a dead cell indicator. Three biological replicates are shown and scale bar is 1 mm.

**TABLE S1:** Values of model parameters used to match experiments.

| Parameter | Units | Values | Reference (or) Rationale for choice |
| --- | --- | --- | --- |
| Maximal growth rate, $g$ | $\text{h}^{-1}$ | 0.32 | This study |
| Maximal death rate, $d$ | $\text{h}^{-1}$ | 0.71 | This study |
| Nutrient consumption rate per cell $\kappa$ | $(\text{mM})(\text{CFU}/\text{mL})^{-1}(\text{h})^{-1}$ | $1.1 \times 10^{-9}$ | This study |
| Critical nutrient concentration, $c_*$ | mM | 0.02 | [9] |
| Critical antibiotic concentration, $a_*$ | $\mu\text{g mL}^{-1}$ | 7 | This study |
| Nutrient diffusivity, $D_c$ | $\text{mm}^2 \text{h}^{-1}$ | 2.2 | [5] |
| Antibiotic diffusivity, $D_a$ | $\text{mm}^2 \text{h}^{-1}$ | 2.2 | $D_a = D_c$ since both are similarly sized small molecules |
| Growth Threshold, $\Phi$ | — | 1.5 | This study |
| Initial bacteria concentration, $b_0$ | $\text{CFU mL}^{-1}$ | $10^8 - 10^9$ | This study |
| Initial glucose concentration, $c_0$ | mM | 0.22 – 2.2 | This study |
| Initial fosfomycin concentration, $a_0$ | $\mu\text{g mL}^{-1}$ | 256 – 2048 | This study |

**TABLE S2:** Ranges of values of model parameters explored in this study with corresponding references.

| Parameter | Units | Range | References (or) Rationale for choice |
| --- | --- | --- | --- |
| Maximal growth rate, $g$ | $\text{h}^{-1}$ | 0.06 – 2 | [19–22] |
| Maximal death rate, $d$ | $\text{h}^{-1}$ | 0.1 – 4 | Death rate scales linearly with growth rate [21] |
| Nutrient consumption rate per cell $\kappa$ | $(\text{mM})(\text{CFU}/\text{mL})^{-1}(\text{h})^{-1}$ | $1 \times 10^{-9} - 5 \times 10^{-8}$ | [6, 23] |
| Critical nutrient concentration, $c_*$ | mM | $2 \times 10^{-4} - 0.5$ | [9, 22] |
| Critical antibiotic concentration, $a_*$ | $\mu\text{g mL}^{-1}$ | 0.1 – 512 | Approximated by MIC [24] |
| Nutrient diffusivity, $D_c$ | $\text{mm}^2 \text{h}^{-1}$ | 0.3 – 3 | [25–27] |
| Antibiotic diffusivity, $D_a$ | $\text{mm}^2 \text{h}^{-1}$ | 0.01 – 2 | [28, 29] |
| Growth Threshold, $\Phi$ | — | 1.5 | This study |
| Initial bacteria concentration, $b_0$ | $\text{CFU mL}^{-1}$ | $4 \times 10^4 - 2 \times 10^{11}$ | [6, 30–32] |
| Initial glucose concentration, $c_0$ | mM | 0 – 48 | [33–35] |
| Initial fosfomycin concentration, $a_0$ | $\mu\text{g mL}^{-1}$ | 0 – 1500 | [36, 37] |

---

**FIG. S1:**
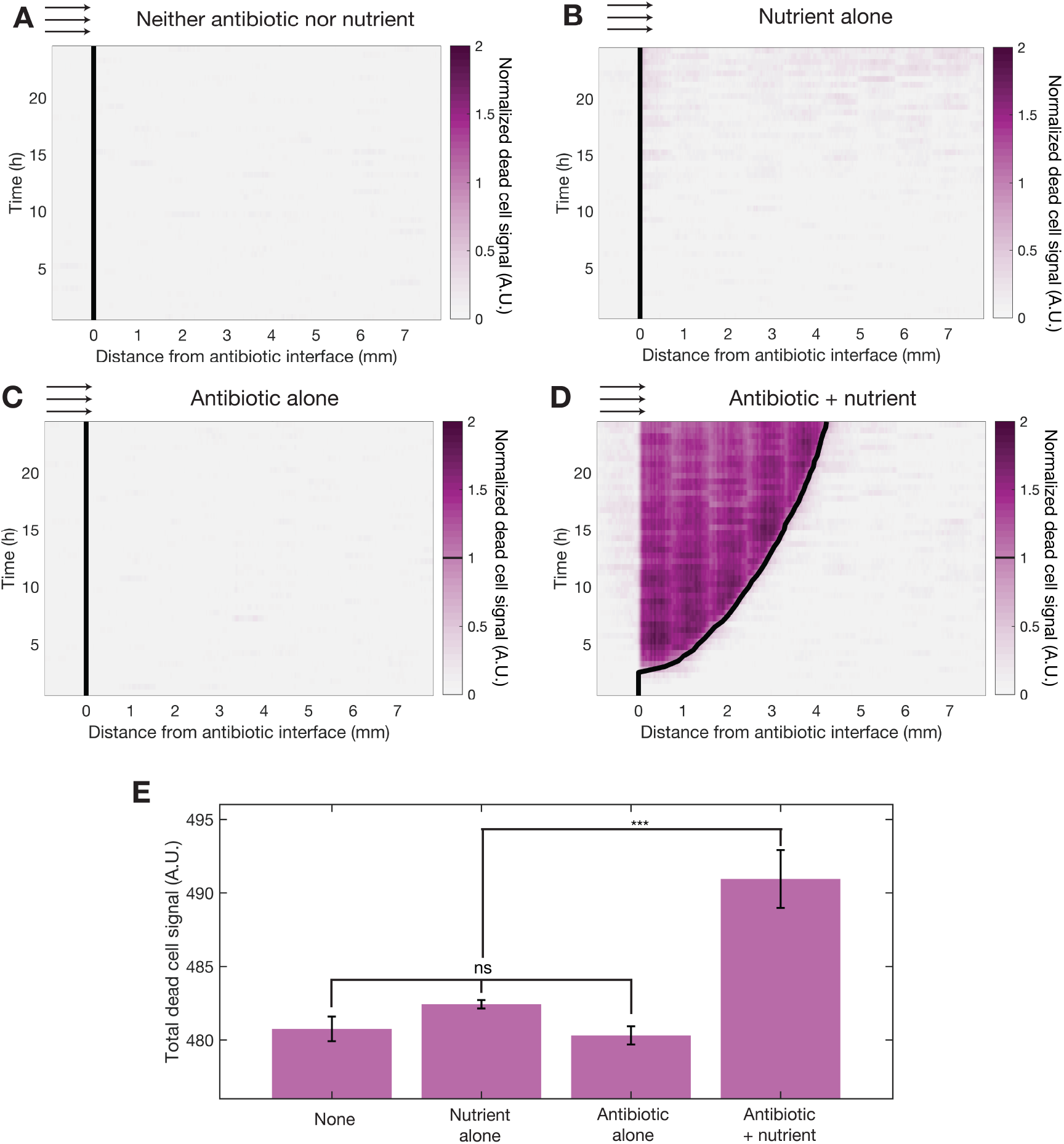
Propidium iodide (PI) dead-cell fluorescent signal from 1 replicate of time-lapse confocal microscopy across conditions. For each condition, normalized dead cell signal is obtained by subtracting a baseline noise value and then dividing by an arbitrary threshold value, both of which are constant across conditions. Then, death front position (black line) is defined as the furthest *x* position with a normalized signal value of 1. **A** Kymograph of PI signal for 10^8^ CFU/mL stationary phase *E. coli* population. No nutrients or antibiotic are placed in the reservoir. **B** Kymograph of PI signal for 10^8^ CFU/mL stationary phase *E. coli* population encountering a diffusing source of 22.2 mM glucose and no antibiotic. **C** Kymograph of PI signal for 10^8^ CFU/mL stationary phase *E. coli* population encountering a diffusing source of 2048 µg/mL Fos-Na and no glucose, the same condition seen in Main Text Fig. 1C and Movie S1. **D** Kymograph of PI signal for 10^8^ CFU/mL stationary phase *E. coli* population encountering a diffusing source of 2048 µg/mL Fos-Na and 0.22 mM glucose, the same condition seen in Fig. 1D and Movie S2. **D** Total dead cell fluorescence signal for each condition **A-D** averaged across 3 biological replicates with standard deviation shown by error bars. Statistical significance is assessed using a 1-way ANOVA test; *** indicates *p <* 0.001 and ns indicates a difference that is not significant. Thus, antibiotic alone does not cause appreciable cell death compared to nutrient rich or nutrient poor environments without antibiotic.

**FIG. S2:**
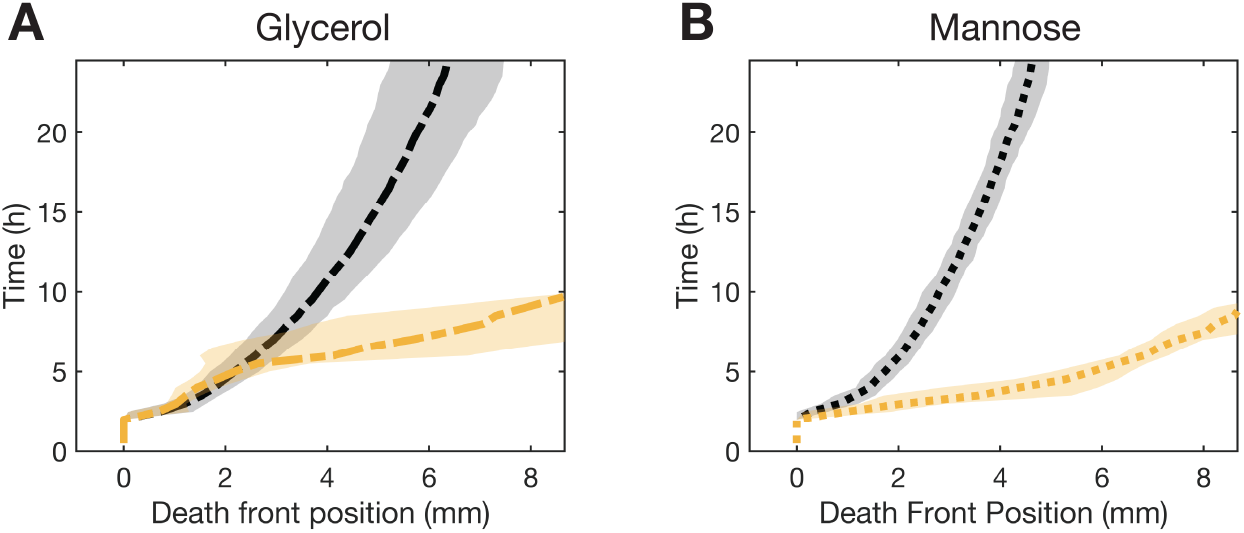
Increasing nutrient source concentration from 0.22 mM (black line) to 2.2 mM (yellow line) **A** glycerol and **B** mannose increases death front clearance speed for a 10^8^ CFU/mL *E. coli* population treated with 2048 µg/mL Fos-Na. Shading around all lines represents standard deviation in death front position at each time point across 3 biological replicates.

**FIG. S3:**
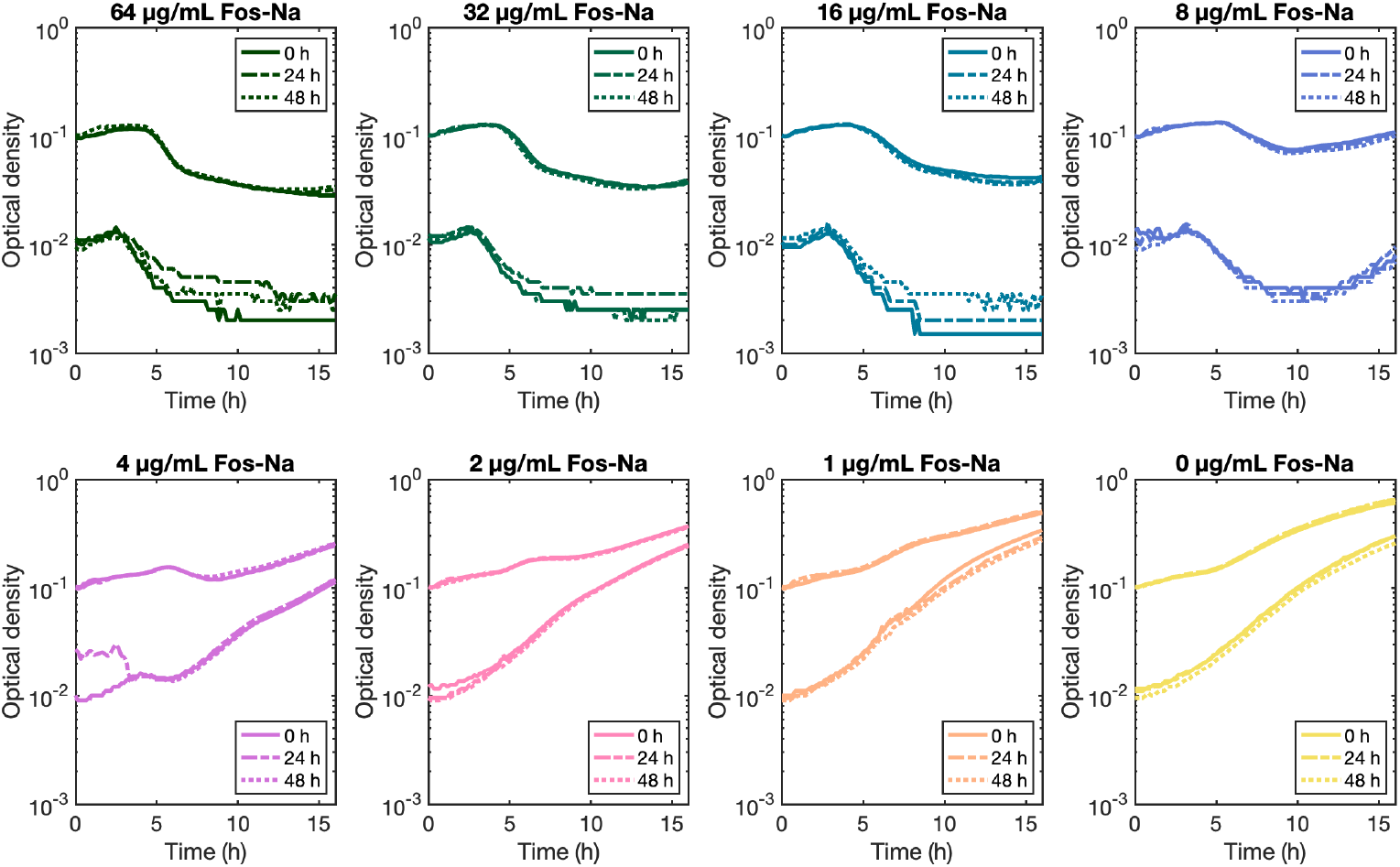
Growth and death curves of cells with fresh vs. pre-incubated antibiotic. Growth and death does not vary across conditions suggesting that fosfomycin does not degrade significantly over the time scale of experiments. Legend indicates time antibiotic stock was pre-incubated at 37°C prior to the experiment. Each line represents mean growth of two replicates.

**FIG. S4:**
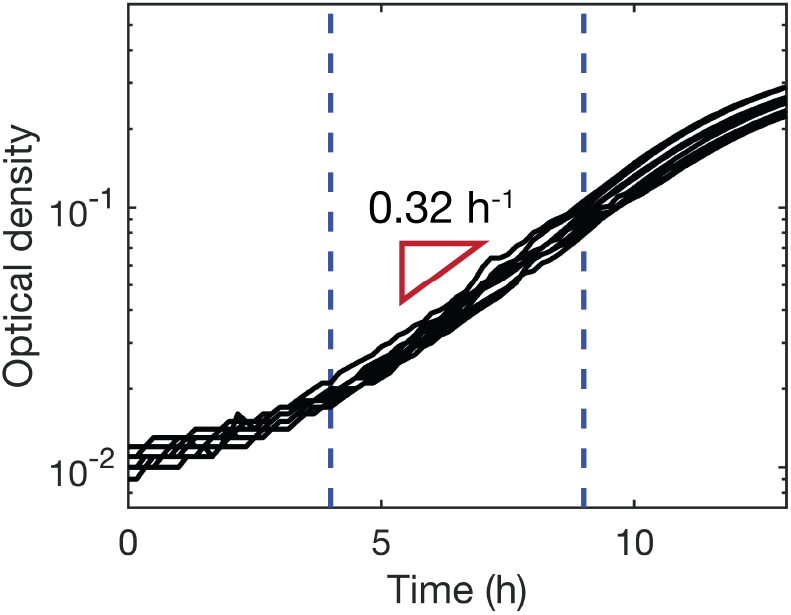
Growth of *E. coli* in 20.0 mM glucose M9 media. Black lines show 8 replicates across 4 independent experiments, each of which was fit with an exponential curve between 4-9 h (blue dotted lines), to yield an average exponential growth rate of 0.32 h^−1^.

**FIG. S5:**
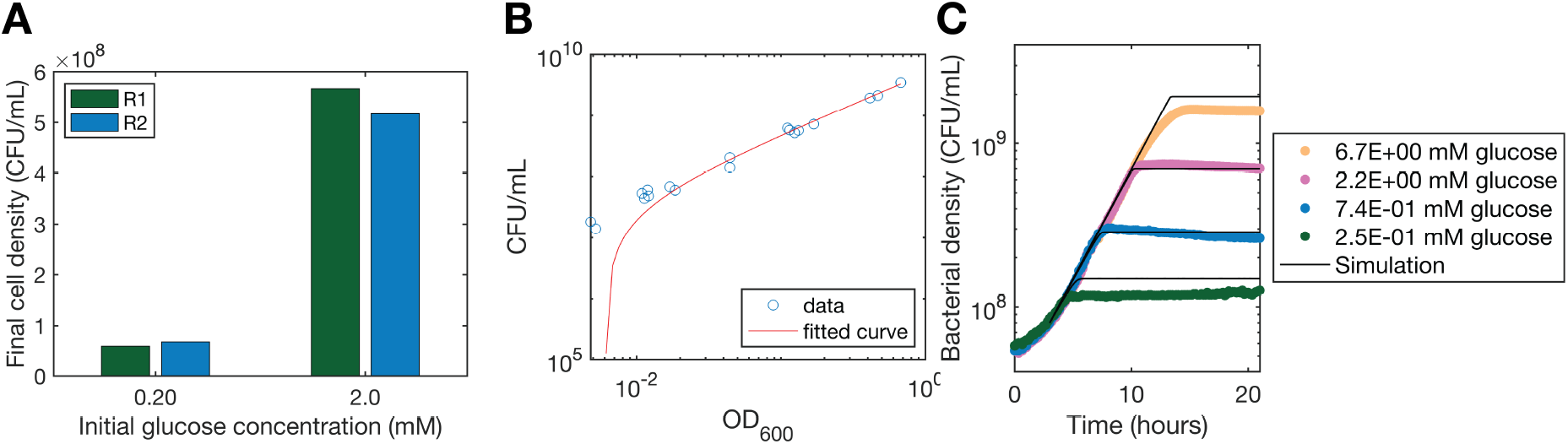
Experiments to parameterize *κ*. **A** Final cell density of two replicate cultures inoculated with *<* 10^7^ CFU/mL. Final yield was used to calculate the nutrient consumption rate per cell, *κ* according to the expression 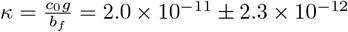 (mM)(CFU/mL)^−1^(h)^−1^. **B** OD_600_ to CFU/mL linear fit conversion from measurements across many independent experiments. **C** Good agreement between experimental growth curves (dots) and well-mixed simulations (black lines) across 4 nutrient conditions confirms an appropriate value for *κ*. Experimental measurements were converted from OD_600_ to CFU/mL according to the linear fit in **B**.

**FIG. S6:**
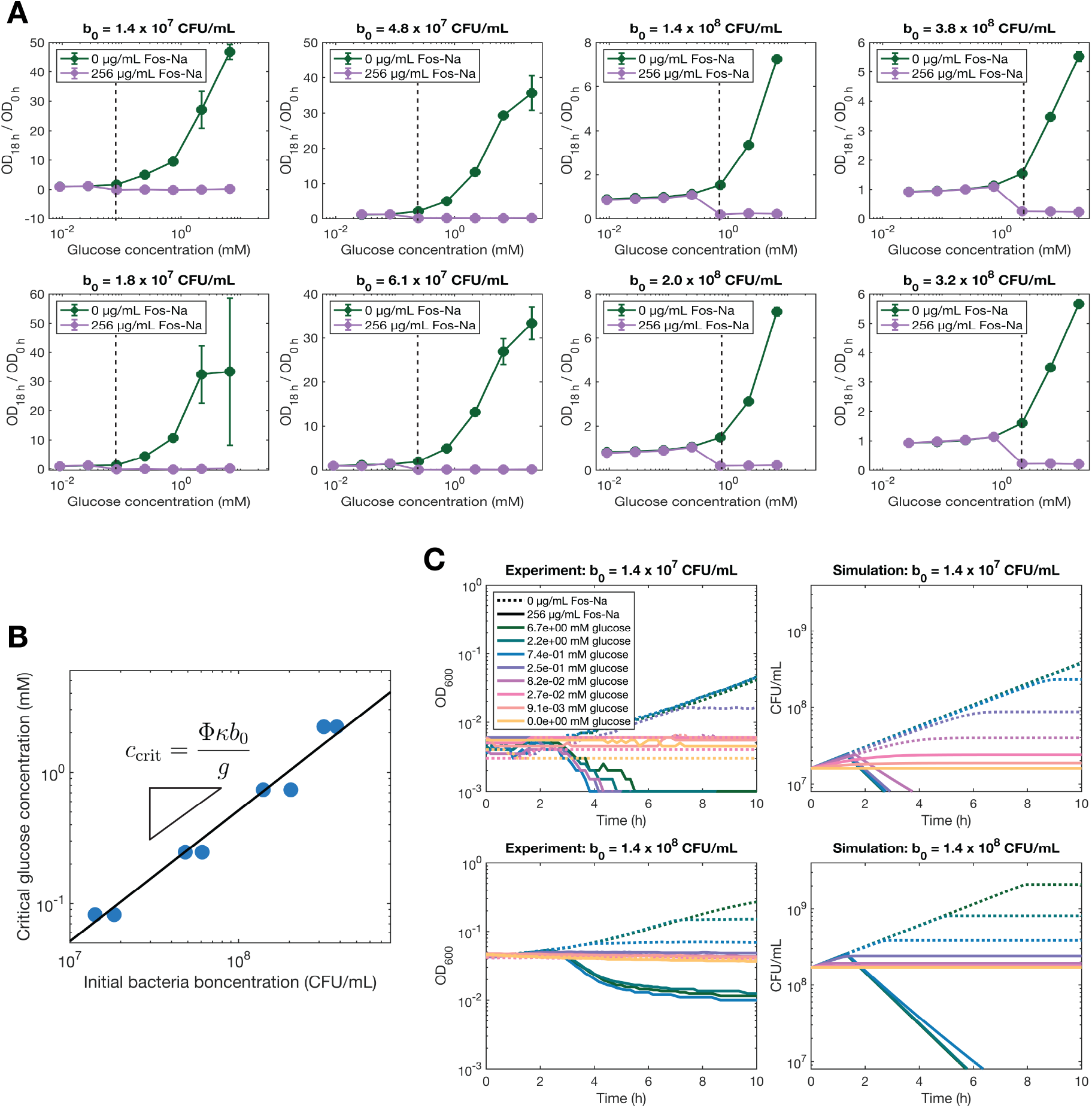
Experiments to parameterize growth threshold parameter Φ. **A** Optical density after 18 h growth compared to initial optical density for varying inoculum densities cultured in a range of glucose concentrations. At and above a threshold nutrient concentration, *c*_crit_, marked by a dashed line on each plot, cells died in the presence of 256 µg/mL Fos-Na. Each point is a mean of two technical replicates with error bars to indicate the standard deviation. Example time course data for two experiments in **A** are shown in panel **C. B** The critical glucose concentration *c*_crit_ varies linearly with initial cell density *b*_0_ as expected from our theory. Each blue dot corresponds to an independent experiment shown in **A** and the black line follows the equation 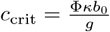, where Φ = 1.5 *±* 0.3 averaged across all 8 experiments. **C** Simulating well-mixed growth with the measured value of Φ gives qualitatively similar time course results to experiments.

**FIG. S7:**
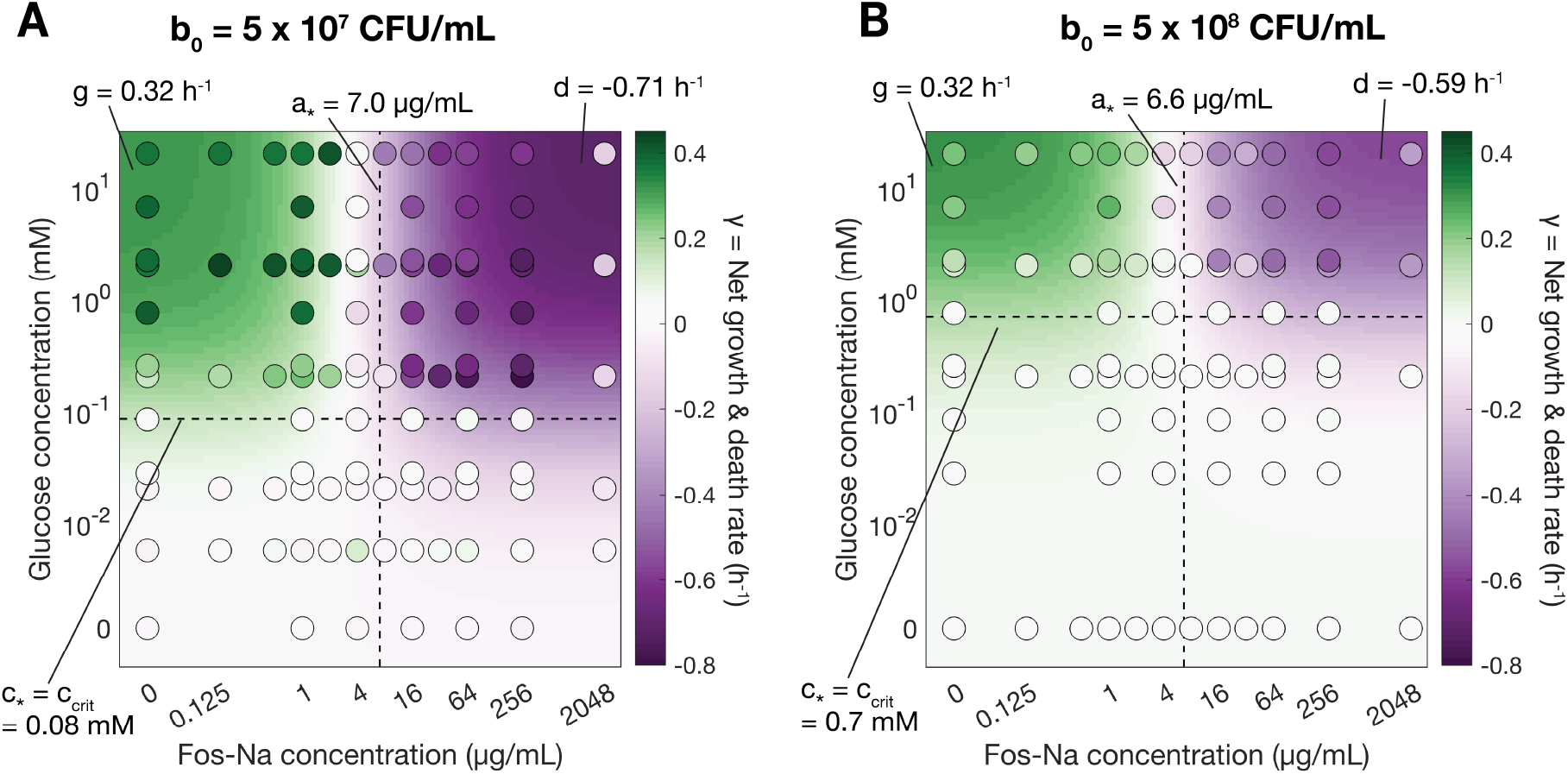
Experiments to parameterize *a*_∗_ and *d*. We measure net cell growth and death rate *γ* as a function of nutrient and antibiotic concentration for **A** low and **B** high initial cell densities. Dots are experimentally measured rates from UV-vis experiments across nutrient, antibiotic, and cell density conditions. Data represents over 100 conditions and each dot is the mean of 2-4 replicates. Background color is the fit for the function 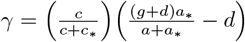 where *g* is fixed to 0.32 h^−1^. Simulations use fit parameters from the low cell density conditions **A**, as summarized in Table S1.

**FIG. S8:**
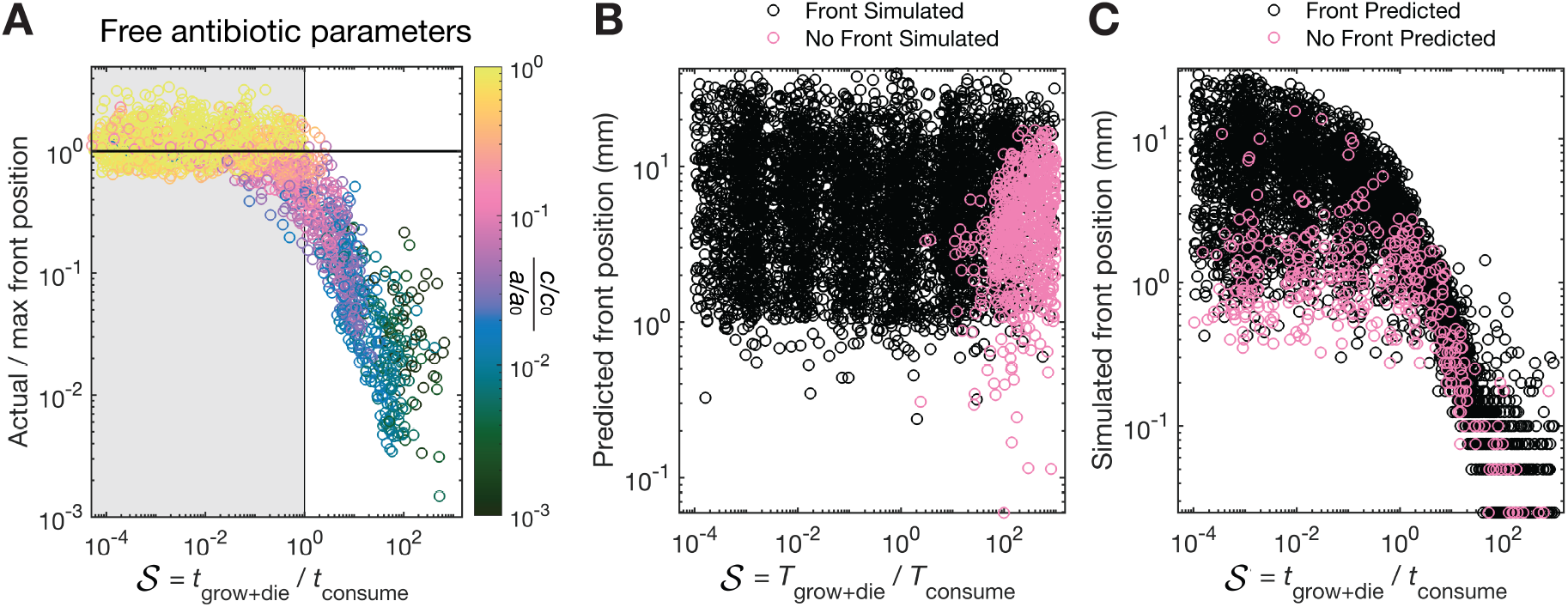
**A** Sweeping across all parameter ranges, without constraining *D*_*c*_ = *D*_*a*_ and *c*_0_*/c*_∗_ = *a*_0_*/a*_∗_, as in Main Text Fig. 4C, the actual simulated front position still collapses away from the theoretical maximum value when the timescale for growth and death *t*_grow+die_ is much larger (slower) than the timescale for nutrient consumption *t*_consume_. Each simulation is evaluated 12 h after a death front is detected and cases where no front forms in the predicted time window are excluded. Point coloring describes the normalized nutrient concentration at the front position vs. the normalized antibiotic concentration at the front. **B** For certain simulations across all presented conditions, no front is simulated (pink points) even when one is predicted. These are concentrated to large 𝒮 and can occur across the range of all predicted front positions (black points). **C** Similarly, for some simulation conditions, no front is predicted to form (pink points) yet a front is simulated. These outlier cases occur across all values of *S* and represent a small fraction of total simulations where a front is successfully predicted.

**FIG. S9:**
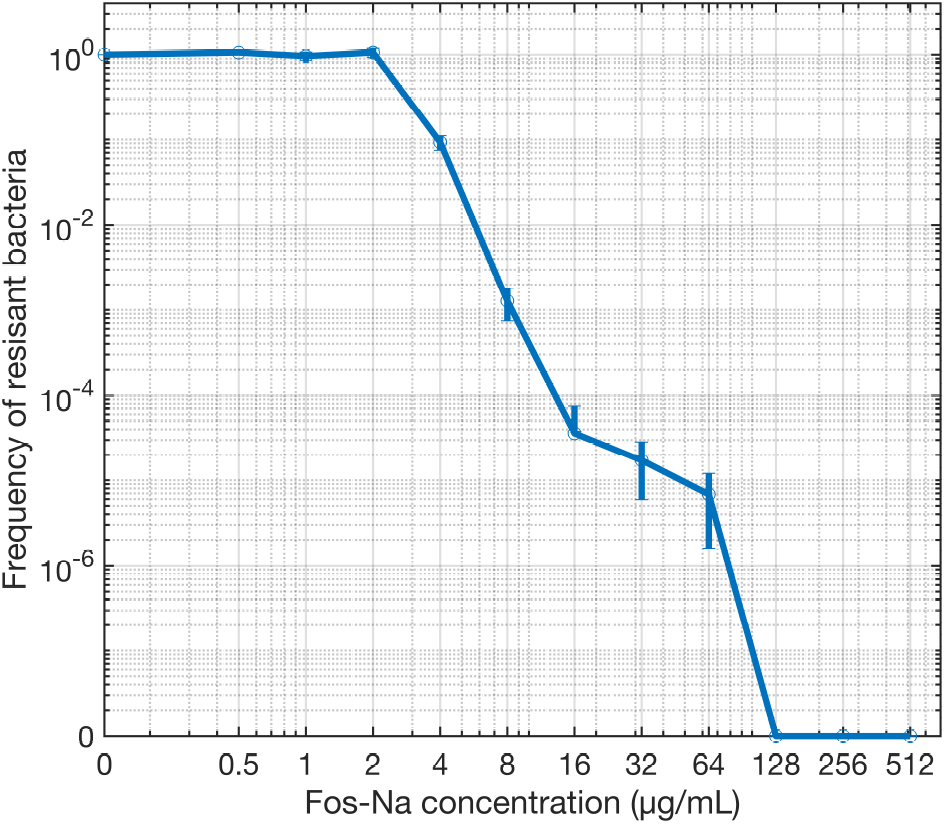
Population analysis profiling of cell strain used in this paper reveals heteroresistance to fosfomycin. CFU counts for cells that grew on at each antibiotic concentration are compared to antibiotic-free CFU counts to obtain the frequency of resistant bacteria. Since resistant subpopulations with frequencies greater than 10^−7^ are detected over an 8-fold concentration range of fosfomycin our cells can be defined as heteroresistant [3]. The frequency detection threshold for the assay is 10^−8^ and antibiotic concentrations where no cells are detected are marked with a frequency of zero. Each dot represents the mean of 3 biological replicates with standard deviation shown as error bars.

**FIG. S10:**
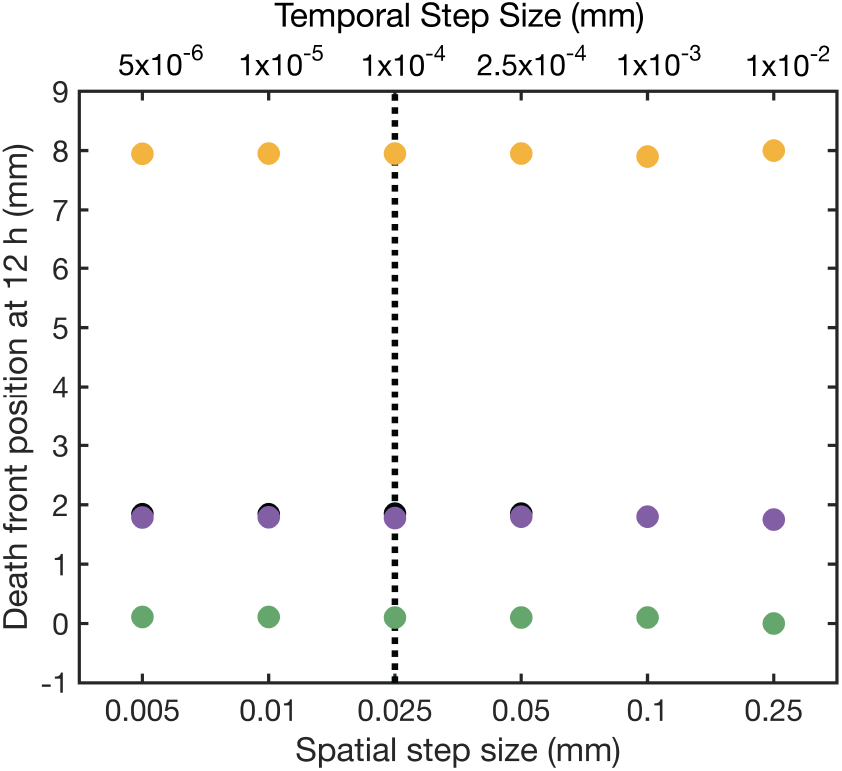
Simulations results do not change appreciably for varying varying spatial and temporal discretization. To assess the sensitivity of our results to numerical discretization, we repeat four representative simulations at varying discretizations. Each color dot represents a different set of initial conditions for the representative simulations. Black is the base case with *b*_0_ = 10^8^ CFU/mL, *c*_0_ = 2.22 mM, *a*_0_ = 2048 µg/mL Fos-Na. Purple has less antibiotic with *a*_0_ = 256 µg/mL Fos-Na. Yellow has more nutrient with *c*_0_ = 0.22 mM glucose. Green has more cells with *b*_0_ = 10^9^ CFU/mL. Dashed line denotes discretization used in the numerical simulations across the paper. The death front position after 12 h obtained from the simulations is not strongly sensitive to the choice of numerical discretization for spatial and temporal steps smaller than those used for the simulations presented in the main text (dashed line). Thus, our choice of discretization is sufficiently finely resolved such that the results in the numerical simulations are not appreciably influenced by discretization.

## Notes

http://doi.org/10.5281/zenodo.14990205

### Competing Interest Statement

The experimental platform used is the subject of a patent application filed by Princeton University on behalf of S.S.D. (PCT/US/2020/030213). All the other authors do not have competing interests.

